# Validating the Gap-Startle Paradigm for Tinnitus Detection: A Machine Learning Approach in CBA/CaJ Mice

**DOI:** 10.64898/2026.09.05.747694

**Authors:** Alyssa B. Price, Dimitri L. Brunelle, Collin R. Park, Andrea S. Lowe, Edward Lobarinas, Joseph P. Walton

## Abstract

Tinnitus is one of the most common hearing disorders affecting one-third of Americans and is defined as the buzzing or ringing sound one perceives in one or both ears in the absence of an acoustic stimulus. One objective method for tinnitus screening in rodents is gap prepulse inhibition of the acoustic startle reflex (GPIAS), a reduction in the abrupt motor response elicited by an intense auditory stimulus following a silent gap in noise. Reduced inhibition by gaps embedded in narrow-band noise is hypothesized to reflect the primary tinnitus pitch, as the tinnitus “fills-in” the gap. However, Lobarinas et al. (2013) identified a critical limitation: after acoustic trauma or hearing loss, rodents often exhibit markedly diminished acoustic startle reflexes, creating a “floor effect” where further suppression becomes undetectable even if gap perception remains intact, leading to false-positive tinnitus screening results. To address this limitation, we utilized the CBA/CaJ mouse model to assess tactile airpuff modification efficacy and employed a machine learning algorithm to classify startle responses, achieving 98% accuracy in differentiating startles from non-startles. We induced unilateral conductive hearing loss via ear plugging and found enhanced gap detection ability, contrasting with false-positive tinnitus indicators observed in rats by Lobarinas and colleagues. We also pharmacologically induced tinnitus via sodium salicylate, revealing frequency-specific alterations in gap detection patterns. Our findings suggest that differences in data analysis methodology, specifically using a machine learning algorithm to filter non-startle responses, may explain species-specific differences between mouse and rat models and significantly improve GPIAS validity as a tinnitus screening assessment tool.

**Highlights:** • Gap-startle paradigm objectively screens for tinnitus in rodent models.

• Machine learning classification achieved 98% accuracy in startle detection.

• Enhanced analysis resolved floor effects that previously limited paradigm validity.

• Species differences explained by improved data filtering methodology.

## 1. Introduction

Subjective tinnitus refers to the perception of ringing or buzzing in the absence of an external acoustic stimulus (De Ridder et al., 2021; Henry & Meikle, 2000). Globally, approximately 14% of adults experience tinnitus, and about 2% report a severe form (Jarach et al., 2022). Despite its prevalence, there are currently no widely adopted or FDA-approved treatments that fully eliminate tinnitus. Populations at increased risk include older adults, military personnel, industrial workers, and musicians (Eggermont & Roberts, 2015). For many individuals, tinnitus is profoundly debilitating, adversely affecting sleep, concentration, emotional well-being, and perceived quality of life (Watts et al., 2018). Consequently, elucidating the neural and behavioral mechanisms underlying tinnitus has significant clinical relevance. Animal models have been instrumental in advancing our understanding of the pathophysiological processes associated with tinnitus (Yang et al., 2007). Among these, the CBA/CaJ (CBA) mouse is frequently employed for tinnitus research because its auditory function and age-related hearing loss patterns closely parallel those observed in humans (Gao et al., 2004).

Tinnitus frequently co-occurs with hyperacusis, a condition characterized by abnormally reduced tolerance to everyday sounds, with estimates suggesting that 30–50% of tinnitus sufferers also experience hyperacusis (Schecklmann et al., 2014; Refat et al., 2021). Both conditions are thought to arise from maladaptive central gain increases following peripheral auditory damage, suggesting shared pathophysiological mechanisms (Eggermont & Roberts, 2015). This comorbidity is particularly relevant in pharmacological animal models of tinnitus, where agents such as sodium salicylate (SS) are known to induce both tinnitus-like percepts and heightened auditory sensitivity concurrently (Chen et al., 2014; Radziwon et al., 2017). To detect these auditory perceptual changes in animal models, researchers have relied heavily on reflex-based behavioral measures. The acoustic startle response (ASR) is a rapid, reflexive motor reaction to a sudden and intense acoustic stimulus, and is a fundamental defensive behavior observed across mammalian species including humans (Davis, 1984; Dawson et al., 1999). For almost a century, the ASR has served as a robust behavioral measure for assessing sensorimotor processing and reactivity to acoustic startle elicitor stimuli (SES) (Landis & Hunt, 1939).

The magnitude of the ASR is attenuated when a brief, non-startling stimulus - known as a prepulse - precedes the SES, a phenomenon termed prepulse inhibition (PPI) (Ison et al., 1973). PPI reflects the brain’s capacity to filter and integrate sensory information over short time intervals and is widely used to assess auditory temporal processing in rodents (Galazyuk & Hébert, 2015). A specialized form of PPI, the gap-in-noise test, employs a brief silent gap occurring within continuous background noise as the prepulse. Under normal hearing conditions, this silent gap suppresses the startle response. However, in animals with tinnitus, the perception of the phantom sound is thought to “fill in” the gap, thereby reducing or abolishing gap-induced startle suppression (Longenecker & Galazyuk, 2011). Consequently, gap-induced PPI has become a widely used behavioral assay for assessing temporal processing and detecting tinnitus in rodents (Barsz & Walton, 2002; Ison et al., 1991; Lobarinas et al., 2013), and has undergone continued methodological refinement to improve its reliability as a screening tool (Longenecker & Galazyuk, 2012). This method is particularly effective in CBA mice, whose auditory sensitivity and temporal processing characteristics make them a well-suited model for tinnitus research.

Turner et al. (2006) developed a tinnitus-screening paradigm that incorporated various narrowband noise carriers containing silent gaps. As a result, the gap would serve as a less effective prepulse, leading to reduced PPI of the ASR. For the GPIAS paradigm to reliably indicate tinnitus, two key conditions must be met: the animal must (1) detect the continuous background noise containing the gap and (2) produce a measurable startle response to the acoustic SES. Longenecker and Galazyuk (2011) demonstrated that unilateral noise exposure in mice reduced ASR amplitude by approximately 52%, an effect that persisted for over three months. Similarly, Lobarinas et al. (2013) reported that temporary unilateral hearing loss induced by earplugging in rats led to apparent tinnitus-like behavior - suggesting that peripheral hearing loss alone can generate false-positive outcomes in GPIAS assays. In these studies, tinnitus-like behavior was quantified using the Gap:No-Gap Ratio (G:NG), which compares the peak startle amplitudes between trials containing a gap (G) and those without (NG). An increased G:NG indicates impaired gap detection and, by inference, a tinnitus-like percept. (Note that PPI and G:NG have a reciprocal relationship.) Earplugging reduced startle amplitudes in both gap and NG conditions but produced an overall increase in G:NG when an acoustic SES was used. In contrast, when a tactile SES (a rapid airpuff) replaced the acoustic startle stimulus, startle amplitudes decreased across conditions, but G:NG values remained unchanged. Thus, unilateral earplugging mimicked a tinnitus-like increase in G:NG under acoustic, but not tactile, startle conditions. Whether this floor effect is specific to rats or also extends to mice, a species increasingly used in tinnitus research, remains unknown.

The present study sought to replicate and extend the findings of Lobarinas et al. (2013) in young adult CBA mice. Our goals were to examine potential species differences between rats and mice, to evaluate the utility of a tactile SES as an alternative readout, and to assess drug-induced tinnitus using sodium salicylate (SS). Two experiments were conducted: one employing earplugs to induce unilateral conductive hearing loss (CHL), and another using SS to pharmacologically induce tinnitus. Through these approaches, we aimed to clarify the sensitivity of GPIAS to tinnitus-related changes in auditory processing in mice and to further validate its translational potential as a behavioral screening tool.

## 2. Materials and Methods

### 2.1. Subjects

A total of 16 young adult CBA mice (2-4 months old, 8 males and 8 females) were used in this study. Each animal was used in Experiment 1 and Experiment 2, following a within-subjects design. The mice were housed 2-5 per cage in Seal-Safe Plus GM500 cages (36.9 x 15.6 x 13.2 cm.) connected to a Box110SS Techniplast ventilated cage rack (West Chester, PA). The vivarium ambient noise levels across the frequency spectrum were as follows: 34.5 dB SPL at 2 kHz, 29 dB SPL at 4 kHz, 16 dB SPL at 8 kHz, 2.7 dB SPL at 16 kHz, and 0.4 dB SPL at 20 kHz. These measurements were taken from inside a cage located in the housing rack using a Quest Electronics Model 1800 sound level meter with the Model OB-300 1/3 octave filter set and a GRAS 40AE ½” Prepolarized Free-field microphone. The sound level meter was calibrated prior to recording with a Quest electronics CA-22 Pistonphone Calibrator. The housing cages were maintained at a constant temperature and humidity, using a 12-hour light-dark cycle (8 A.M. to 8 P.M.) at 25 ◦C with *ad lib* access to food and water. Behavioral testing occurred during the light cycle. Animals were closely monitored throughout the study period. All experimental procedures used in the present study were approved by the University of South Florida Institutional Animal Care and Use Committee.

### 2.2. Testing apparatus and general procedures

Behavioral testing experiments were conducted using methods previously utilized in this lab (Halonen et al., 2016; Lowe and Walton, 2015; Brecht et al., 2022; Brunelle et al., 2025). The startle response was measured inside one of four identical sound-attenuating boxes (40 x 40 x 40 cm) lined with high frequency anechoic foam. Mice were individually tested in acoustically transparent, 3D-printed mesh cages (9.5 x 4 x 4 cm) mounted on a 3D-printed platform connected to piezoelectric transducers. Transducer responses to their movement in millivolts (mV) were recorded for a window from 125 ms before to 475 ms after the startle stimulus.

Two types of SES were used in the study, acoustic and tactile. Acoustic stimuli were presented through Fostex model FT17H Horn Super Tweeter speakers (Fostex Company, Tokyo, Japan) located 30 cm directly above the transducer platform and were controlled by a RZ6 multi-I/O processor from Tucker-Davis Technologies (TDT, Alachua, FL). Custom MATLAB (The MathWorks, Inc., Natick, MA) and TDT RPvds software were used to control the hardware. All acoustic signals were calibrated prior to testing with a GRAS 40BE 1/4” Prepolarized Free-field Microphone placed at the level of the animal’s pinna in the ASR chamber; voltage output of the acoustic signal was measured via a Tektronix TDS 2014C oscilloscope (Tektronix, Beaverton, OR) and frequency content was measured via a HP3662 spectrum analyzer (HP, Palo Alto, CA) and led to a Larsen Davis preamplifier (model 2221, PCB Piezotronics, Inc., Depew, NY). The tactile startle reflex was elicited with a 20 ms airpuff (at 15 PSI) (Figure 1) delivered to the back of the neck by triggering a solenoid air valve (Parker 2-way solenoid valve VAC-20) placed 3 cm above the animal in the startle chamber. The tactile stimulus also produced an intense acoustic noise burst, so the tactile stimulus itself produced an acoustic signal. A tactile airpuff was used as the startle stimulus rather than an acoustic stimulus to avoid false-positive tinnitus screening arising from hearing loss-induced suppression of the acoustic startle reflex, a known confound in unilateral CHL and salicylate models (Lobarinas et al., 2013).

**Figure 1.**
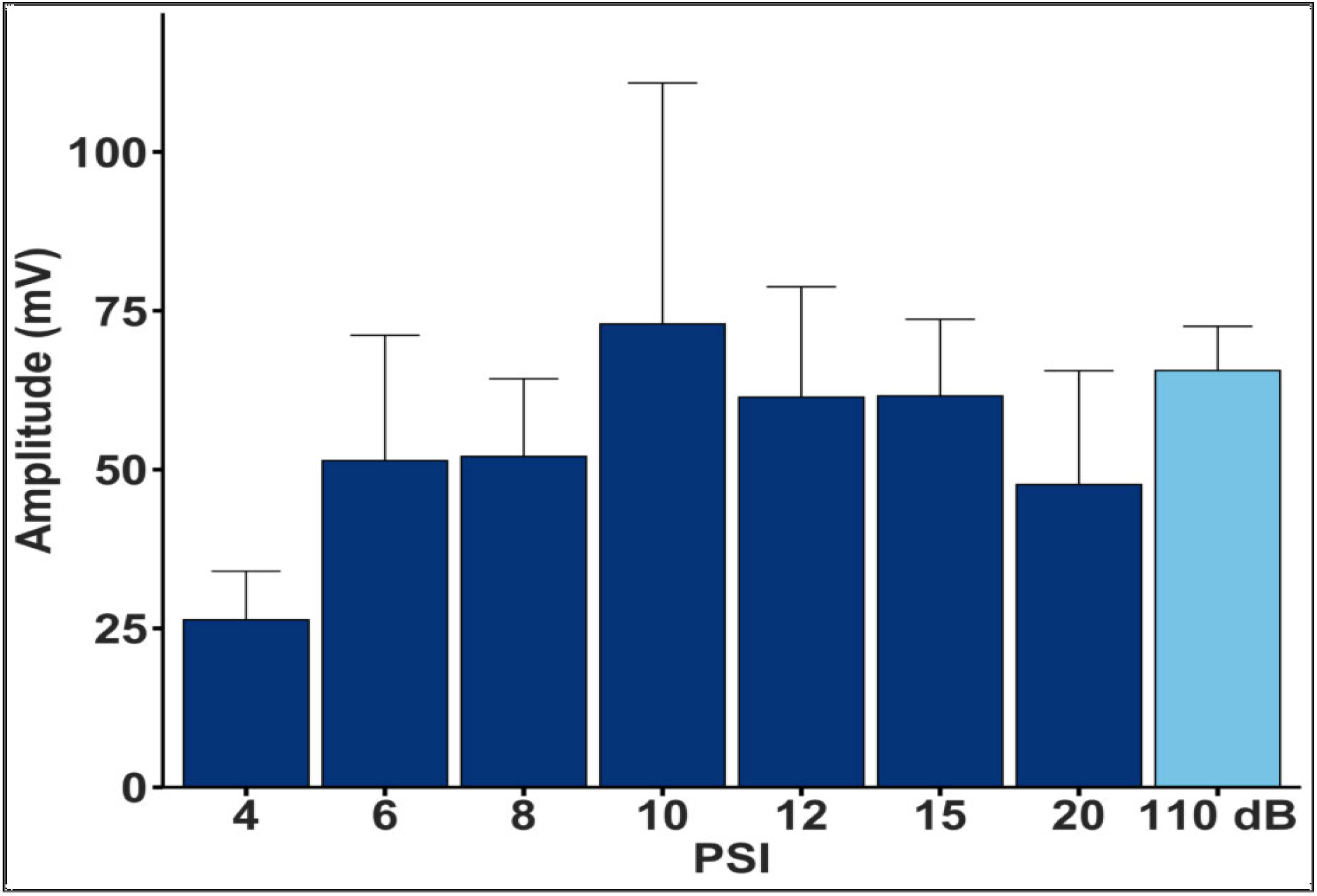
Startle response as a function of air pressure was measured. The optimal air pressure setting for the tactile stimulus was determined to be 15 PSI which yielded the best ratio of amplitude to error (n=8). The lines displayed on each bar indicate SEM.

### 2.3. Startle Data Analysis

#### Startle waveform classification

ASR waveforms were categorized as either startles or non-startles utilizing the *asrclassify* model, a custom machine learning algorithm which has been validated on various rodent models and measurement modalities, developed by our research team in the *R* programming language (Fawcett et al., 2020; 2021; 2023). This model comprises a combination of different machine-learning models that were trained using a dataset of startle waveforms manually classified by several trained auditory neuroscientists. Following feature engineering and training, the model achieved a prediction accuracy of 98% in differentiating startles from non-startles, specifically in CBA mice using piezoelectric transducers. Alternative methods for ASR classification like template matching, thresholding, root mean square (RMS) calculations (Grimsley et al., 2015), and the invalid-trials approach (Schilling et al., 2017) are susceptible to high false-positive rates, leading to significant inaccuracies in mean amplitude calculations and, consequently, generating imprecise experimental vs. control condition amplitude ratios (e.g., G:NG metrics). Therefore, the utilization of the *asrclassify* machine learning algorithm for classifying startle waveforms resulted in a high-quality dataset for subsequent ASR analysis, with non-startle responses being excluded from further examination.

#### Startle waveform metrics

Startle amplitudes were measured for each startle waveform, then averaged across all sessions per mouse within each group. Startle rates were calculated as the total number of true startles divided by the total number of trials per session, then averaged across all sessions per mouse within each group. Startle amplitudes for true startles were measured in mV as the maximum amplitude during the 375 ms time window following stimulus onset.

### 2.4. Experiment 1 – Conductive Hearing Loss

Along with an audible acoustic stimulus, it is known that mice will exhibit a startle reflex when a pressurized airpuff is delivered to the dorsum and thorax of mice (García-Hernández & Rubio, 2022). Based on previous reports, the traditional gap-startle paradigm was modified by replacing the acoustic startle stimulus with a tactile stimulus. In our first experiment, we induced temporary unilateral CHL by earplugging, a manipulation previously shown to generate false-positive indications of tinnitus in rats (Lobarinas et al., 2013). The earplugging method was utilized to determine whether a reduction in startle reactivity could produce false positive results of tinnitus, as seen in the rat model, given the unlikelihood that CHL will induce tinnitus (Bauer et al., 2001). During the first week, mice underwent baseline behavioral testing followed by auditory brainstem responses (ABRs) testing to assess baseline auditory thresholds. Unilateral CHL was induced two weeks later, and ABRs were immediately performed to assess the level of attenuation produced by the earplug. To induce unilateral CHL, a small cotton otoblock was inserted into the left ear canal of each mouse under light isoflurane anesthesia (1.5–2%) and the canal was filled with a silicone elastomer (Kwik-Sil) via injection. The fast-drying elastomer provided a secure and tightly sealed earplug that could not be removed by the mice as it was deeply inserted into the canal. The attenuation of the elastomer was verified by comparing baseline ABRs to those collected immediately after the earplug was inserted (earplug ABRs). Two weeks following earplug insertion, mice underwent behavioral testing. Note that throughout the manuscript, ‘post-earplug’ refers to conditions when the earplug was in place.

### 2.5. Auditory Brainstem Responses

ABRs were obtained using a Tucker Davis Technologies System 3 Real Time Signal Processing System (Tucker Davis Technologies, Alachua, FL, USA). Mice were anesthetized with ketamine (120 mg/kg) and xylazine (10 mg/kg) via intraperitoneal injection for ABR threshold testing. Body temperature was kept constant at 37◦C using a feedback-controlled heating pad (Physiotemp TCAT2-LV Controlller, Clifton, NJ). Subdermal electrodes were placed at the ipsilateral pinna (reference) and the vertex (active), with an electrode at the contralateral pinna serving as a ground. The contralateral ear (non-silicone plugged) was occluded with a pediatric ear probe filled with plumber’s tack during testing. Tone bursts (2 ms, 0.5 rise-fall time, 21/s) of 6, 10, 16, 20, and 24 kHz at intensities ranging from 0–80 dB SPL were presented from TDT MF1 speakers. ABR evoked potentials averaged over 512 repetitions, amplified (RA16PA, Tucker Davis Technologies), filtered (100 - 3000 Hz bandpass), and digitized. The threshold at each frequency was defined as the lowest sound intensity to generate a visible, reproducible waveform (Lowe et al., 2015). Muscle artifacts exceeding 7uV were rejected from the averaged response. All recordings took place in a sound-proof booth lined with echo-attenuating acoustic foam.

### 2.6. Experiment 2 – Tinnitus Induction

Baseline data was collected for Startle Input/Output (SIO) and gap testing 1-2-days prior to SS being administered. To induce tinnitus, 250 mg/kg SS was administered 4 hours prior to testing (n=16). Injections of SS were administered in the morning around 8 A.M., and the various behavioral test types began at 12 P.M..

### 2.7. Testing Procedures

The behavioral tests used were a 1) SIO function, obtained by varying the amplitude of the acoustic startle elicitor at 50, 70, 80, 100 and 110 dB SPL, with 20 trials at each level in a pseudo-random sequence, and 2) a multi-frequency gap (MFG) detection assessment. Gap detection was assessed using a GPIAS procedure in which a 50 ms silent gap was embedded in a continuous background noise presented at 70 dB SPL, positioned 100 ms (onset to onset) prior to the SES at 110 dB SPL. Background noise carriers were narrow-band noises with a bandwidth of 1000 Hz centered at 10, 16, and 20 kHz, as well as a wide-band noise spanning 500 Hz to 40 kHz. For each noise band, 20 trials were presented with a gap and 20 trials without a gap in a pseudo-random order. Animals underwent four sessions of GPIAS testing over a period of 12 weeks, with sessions separated by 2 days to minimize habituation effects (Figure S1).

### 2.8. Data Analysis and Statistics

For the results to be directly comparable to Lobarinas, we calculated the G:NG from the startle amplitudes in the two conditions. NG represented the startle trials without a preceding gap in noise, and Gap represented the startle trials with a preceding gap in noise. Gap detection was impaired at a particular carrier frequency if there was a statistically significant elevation in the G:NG during conductive hearing loss or drug-induced tinnitus compared to the baseline level. Statistical analyses were conducted in the *R* programming language using the base *R* functions aov for ANOVAs and TukeyHSD for Post-Hoc tests. These functions were used to perform two-way analyses of variance (ANOVA) or one-way ANOVAs, depending on the comparison of interest (see Results section for the details of each specific comparison). All statistical comparisons used an alpha value of 0.05. When an ANOVA showed a significant main effect, post-hoc testing was performed with TukeyHSD tests to avoid type I errors associated with multiple comparisons. All results are presented as mean and SEM.

## 3. Results

### 3.1. Experiment 1: Effect of Unilateral Conductive Hearing Loss on Hearing Thresholds

Earplugging produced significant ABR threshold elevations across all test frequencies [*F*(1, 53) = 108.8, p < .05] (Figure 2). Non-earplugged mice exhibited typical audiometric characteristics for young, normal-hearing CBA mice, with the highest thresholds at 6 kHz (47 dB SPL) and lower thresholds at 10 kHz and above (15 dB SPL). Following earplug insertion, hearing thresholds for frequencies at and above 10 kHz were 27 dB SPL higher on average, confirming significant conductive hearing loss. Interestingly, there was not a significant effect of frequency on hearing thresholds in the earplug condition, as the drop off from 6 to 10 kHz was only 9 dB SPL. A one-way ANOVA revealed no significant differences in hearing threshold across frequencies for the post-earplug condition, suggesting a reduced frequency-encoding specificity through the auditory system of these mice.

**Figure 2.**
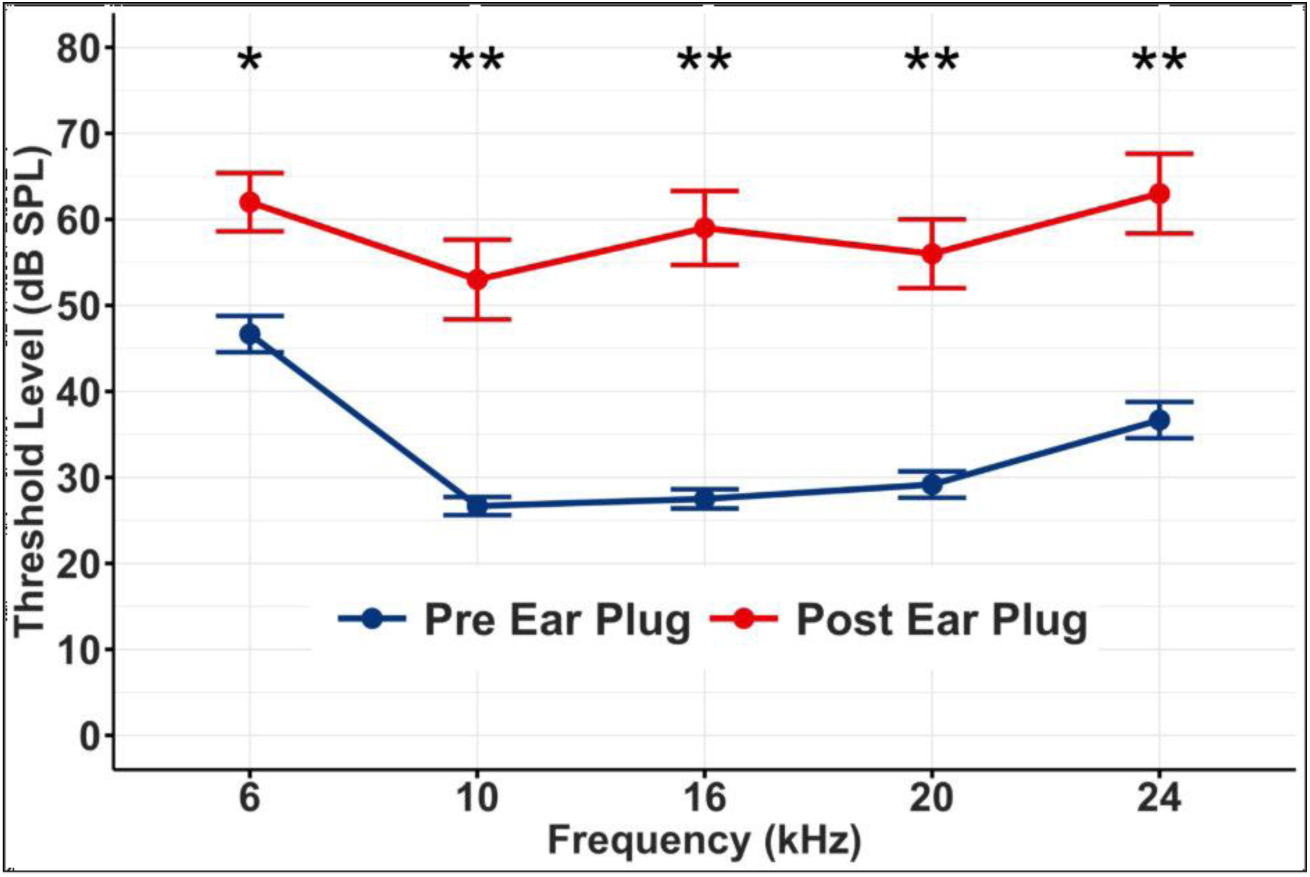
Auditory Brainstem Response (ABR). Hearing thresholds significantly increased following insertion of the ear plug. The conductive loss induced by the earplug ranged from 20-30 dB. The bars displayed on each point indicates SEM and asterisks indicate a significant difference in thresholds between pre-and post-earplug (**\*** P ≤ 0.05, **\*\*** P ≤ 0.01, **\*\*\*** P ≤ 0.001, **\*\*\*\*** P ≤ 0.0001).

### 3.2. Experiment 1: Effect of Unilateral Conductive Hearing Loss on Startle Responses

To assess the impact of conductive hearing loss on startle reactivity, we first investigated startle IO functions using both acoustic and tactile stimuli. For acoustic startle IO testing (Figure 3A), a two-way ANOVA with SES level and earplug condition as factors revealed significant main effects of both level [*F*(1,119) = 18.59, p < .05] and timepoint [*F*(1,119) = 10.48, p < .05], along with a significant interaction [*F*(4,115) = 2.88, p < .05]. Post-hoc testing confirmed significantly decreased startle amplitudes at 100 and 110 dB SPL following earplug insertion. When the tactile startle elicitor was presented (B), a trend toward reduced amplitudes was also revealed [t(28) = -1.92, p = 0.07], likely reflecting attenuation of the acoustic component of the airpuff by the earplug (Plappert et al., 2004; Taylor et al., 1991).

**Figure 3.**
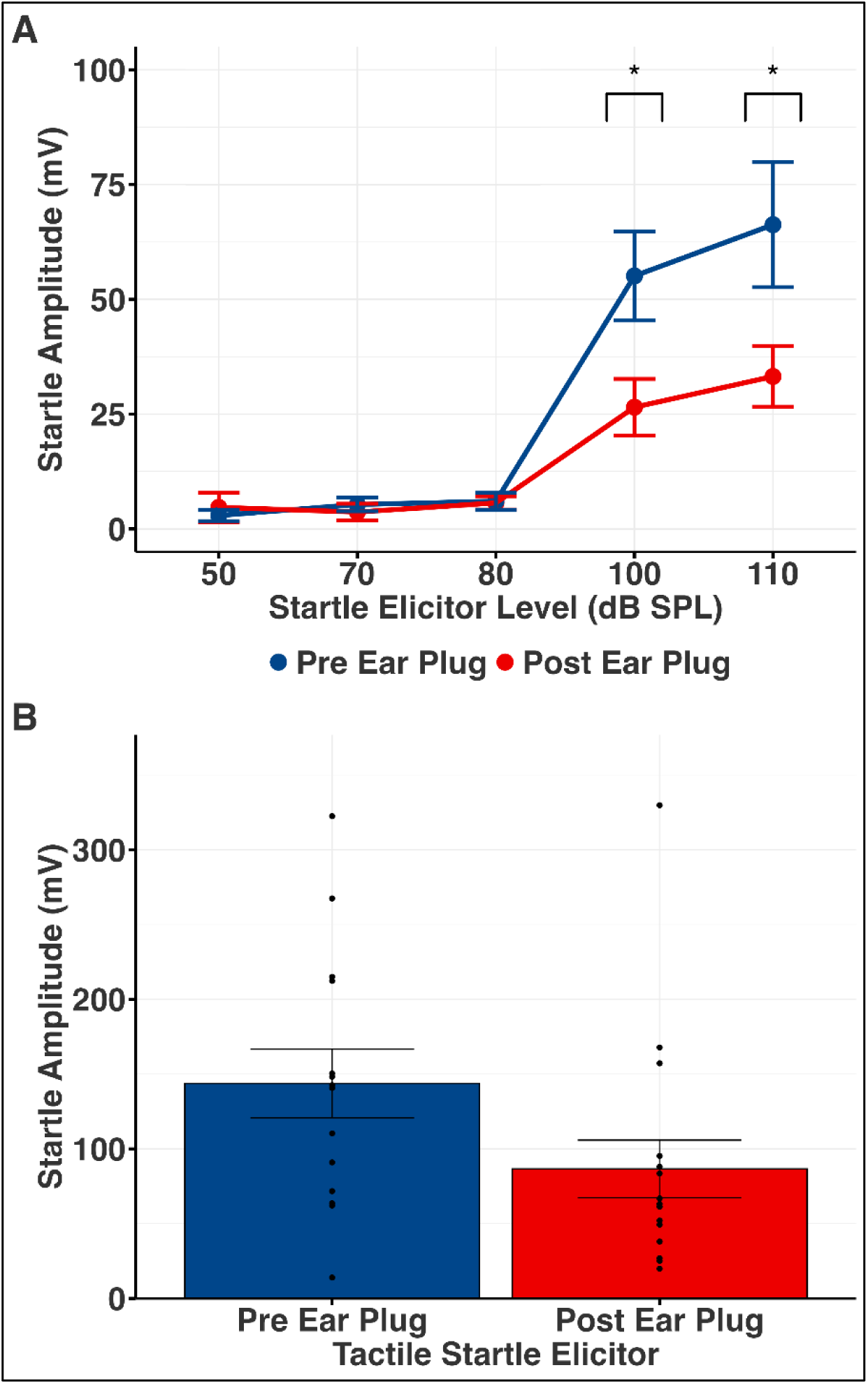
Startle Input/Output function. **A)** The input/output function of the acoustic startle response was decreased in mice post-earplug at SES levels of 100 and 110 dB SPL. **B)** For the tactile SES, there is a decrease in startle amplitudes. This effect was likely due to the acoustic component of the SES being partially eliminated by the earplug. The lines displayed on each point and bar represent SEM and asterisks indicate significant differences between pre- and post-earplug (**\*** P ≤ 0.05, **\*\*** P ≤ 0.01, **\*\*\*** P ≤ 0.001, **\*\*\*\*** P ≤ 0.0001).

We next evaluated gap detection ability using the MFG procedure. When assessing G:NG with an acoustic SES (Figure 4A), a two-way ANOVA identified significant main effects of frequency [*F*(3,119) = 4.56, p < .05] and timepoint [*F*(1,116) = 20.24, p < .05], with no significant interaction [*F*(3,113) = 0.79, p > .05]. Significant decreases in G:NG for the tactile SES were observed for frequency and timepoint [frequency: F(3,113) = 4.2, p < .05; timepoint: F(1,113) = 4.56, p < .05; interaction: F(3,110) = 0.98, p > .05]. Specifically, there were deficits at 16kHz and 20kHz (See post-hoc table, S2).

**Figure 4.**
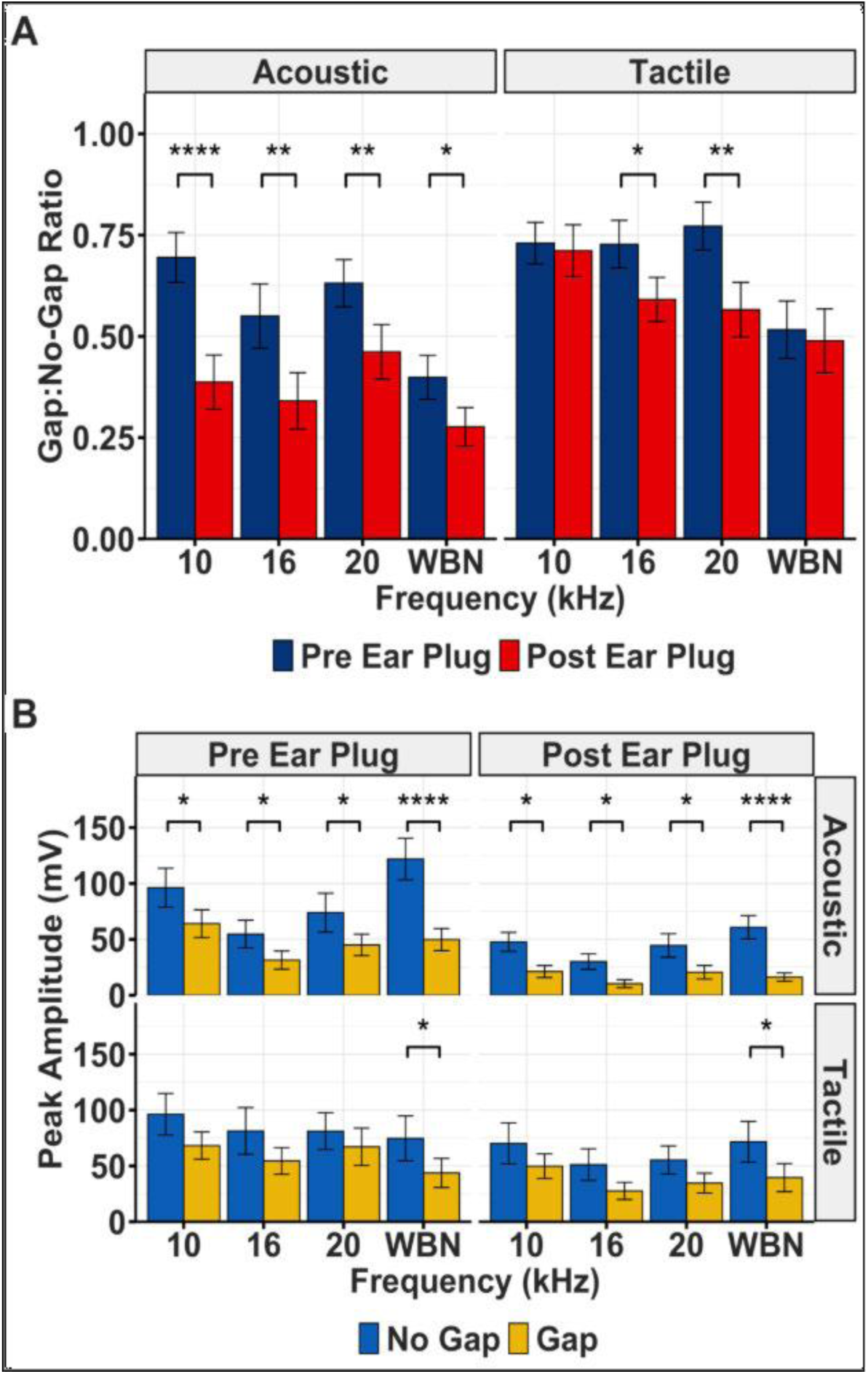
Gap prepulse inhibition of startle. **A)** Using an acoustic startle elicitor, the Gap:No-gap ratio was significantly decreased post-earplug at all carrier noise frequencies. When a tactile startle elicitor was used the Gap:No-Gap ratio was significantly decreased at the 16 and 20 kHz carrier frequencies. **Startle amplitudes.** The lines displayed on bar display SEM and each asterisk indicates significance between gap and no gap trials. **B)** Pre-earplug. Startle amplitudes of the Gap trials were generally lower than No-Gap trials, this difference only reached significance during the acoustic startle with the wide-band noise carrier. **Post-earplug**.

Analysis of peak startle amplitudes for the acoustic SES revealed significant main effects of trial type [*F*(1,239) = 35.17, p < .05] and timepoint [*F*(1,239) = 39.23, p < .05], with no significant interaction [*F*(1,233) = 0.79, p > .05] (Figure 4B). Amplitudes decreased from pre- to post-earplug for both trial types, consistent with a hearing loss-induced reduction in acoustic sensitivity. Importantly, gap trials consistently produced lower amplitudes than NG trials in both earplug conditions, demonstrating preserved PPI. Similar patterns also emerged for the tactile SES [timepoint: *F*(1,233) = 8.04, p < .05; trial type [*F*(1,233) = 11.09, p < .05; interaction: [*F*(1,232) = 0.002, p > .05].

Startle amplitudes of the Gap trials using an acoustic SES were significantly lower than amplitudes of the No-Gap trials, using the 10 kHz, 16 kHz and wide-band carriers. Using a tactile SES, amplitudes during Gap trials were not significantly different than No-Gap trials in either condition. The lines on each bar represent SEM and each asterisk indicates a significant difference between pre- and post-earplug (**\*** P ≤ 0.05, **\*\*** P ≤ 0.01, **\*\*\*** P ≤ 0.001, **\*\*\*\*** P ≤ 0.0001).

### 3.3. Experiment 2 – Pharmacologically Induced Tinnitus via Sodium Salicylate

To establish a positive control for tinnitus detection using GPIAS, we pharmacologically induced tinnitus via injection of SS. SIO testing with the acoustic SES (Figure 5A) revealed significantly elevated amplitudes across all intensities post-SS. A two-way ANOVA confirmed significant main effects of timepoint [*F*(1,151) = 42.20, p < .05] and SES intensity [*F*(4,151) = 44.36, p < .05], along with a significant interaction [*F*(4,147) = 4.94, p < .05]). This hyperresponsiveness likely reflects SS-induced hyperacusis. The tactile SES similarly produced increased amplitudes post-SS (t(30) = 3.62, p = 0.001) presumably due to an enhanced response to the acoustic component of the airpuff (Radziwon et al., 2017).

**Figure 5.**
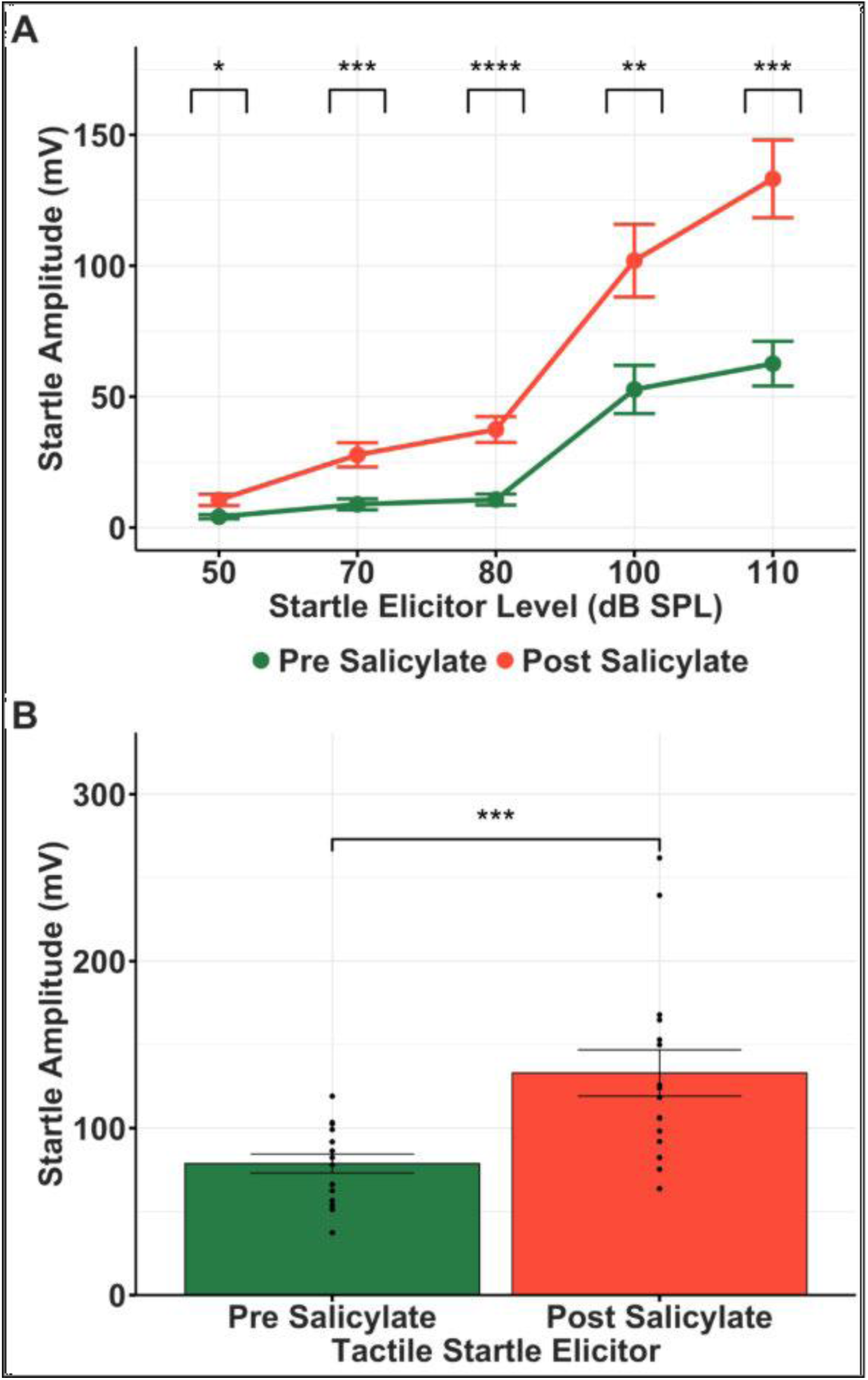
Startle Input/Output Functions. **A)** Using an acoustic SES, startle response amplitude was significantly increased post-SS across all SES levels. **B)** Using a tactile SES, startle response was significantly increased post-SS. The lines displayed on each point or bar represent SEM and each asterisk indicates a significant difference between pre- and post-SS (**\*** P ≤ 0.05, **\*\*** P ≤ 0.01, **\*\*\*** P ≤ 0.001, **\*\*\*\*** P ≤ 0.0001).

Gap detection assessment was then performed pre- and post-SS and revealed frequency-specific alterations in G:NG (Figure 6A). For the acoustic SES, a two-way ANOVA showed a significant main effect of frequency [*F*(3,123) = 21.76, p < .05] but not timepoint [*F*(1,123) = 1.366, p > .05], with a significant interaction [*F*(3,120) = 7.09, p < .05]. This pattern of variable changes across frequencies is consistent with frequency-specific tinnitus percepts. With the tactile SES, the ANOVA revealed a main effect of frequency [*F*(3,123) = 15.41, p < .05] but not timepoint [*F*(1,123) = 0.07, p > .05], with no significant interaction [*F*(3,120) = 1.69, p > .05]. Post-hoc testing revealed that G:NG was significantly higher at 10 kHz relative to 16, 20, and WBN (all p < .05), but no significant differences were found between narrowband frequencies (See post-hoc table).

**Figure 6.**
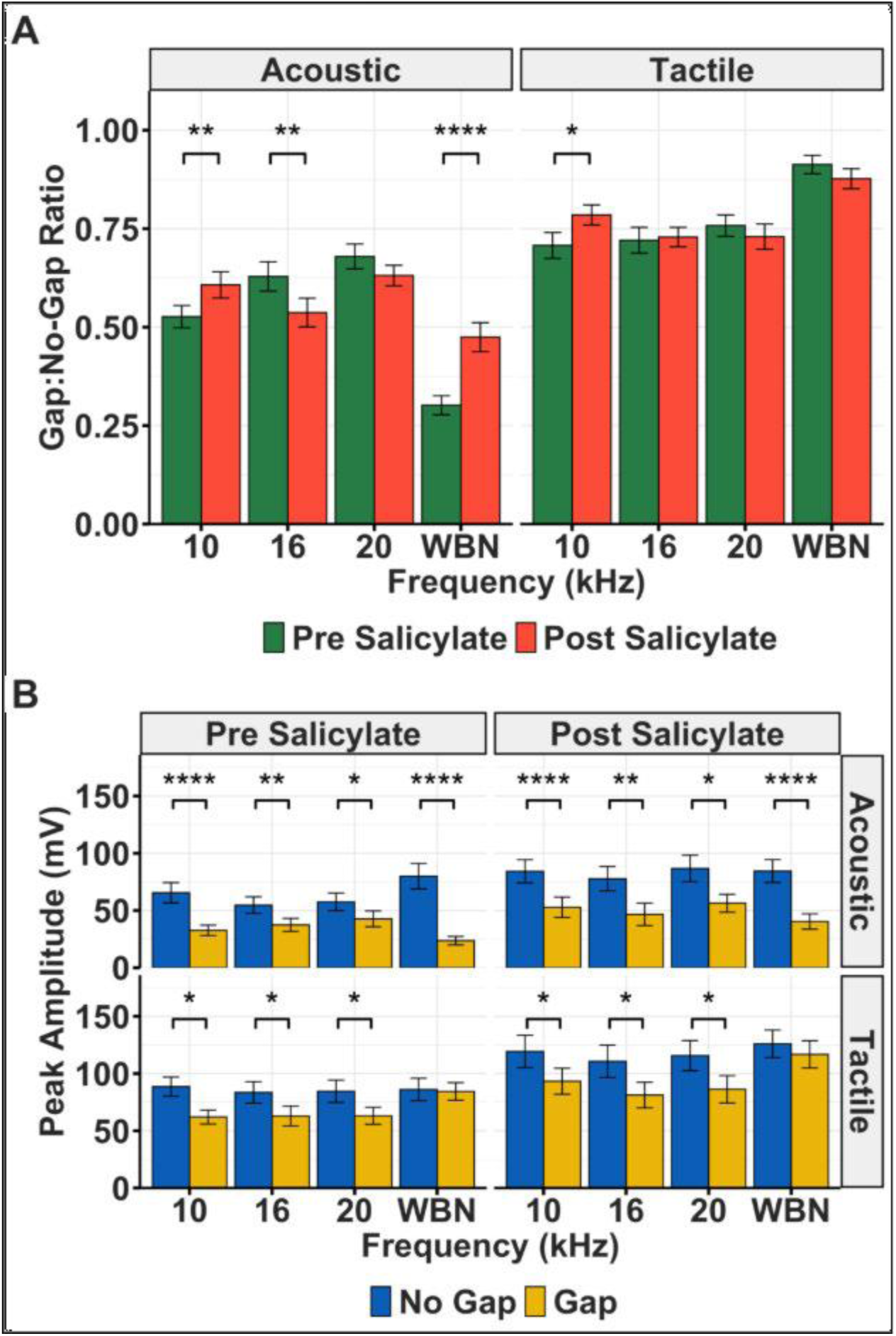
Gap prepulse inhibition of startle. **A)** When evaluating Gap: No-Gap Ratio, in the acoustic startle elicitor stimulus (SES), we see a varying result of significant decrease and increases and non-significant results as well from baseline to treatment at all frequencies. In the tactile SES type, we see a significant increase only at 10k. **Startle Amplitudes. B)** When comparing pre- to post-SS, there is an increase. This effect could be explained by changes in hearing sensitivity that take place from pre- to post-SS due to possible hyperacusis. The effect is also seen when comparing pre- to post-SS for the tactile SES type as well. The lines displayed on each bar represent SEM and each asterisk indicates a significant difference between pre- and post-SS (**\*** P ≤ 0.05, **\*\*** P ≤ 0.01, **\*\*\*** P ≤ 0.001, **\*\*\*\*** P ≤ 0.0001).

Peak startle amplitudes demonstrated consistent increases from pre- to post-SS treatment (Figure 6B). For the acoustic SES, a two-way ANOVA revealed significant main effects of timepoint [*F*(1,253) = 15.90, p < .001] and Gap versus No-Gap trial types [*F*(1,253) = 57.99, p < .001], but not frequency [*F*(3,251) = 0.37, p > .05], with no significant interaction [*F*(3,248) = 0.25, p > .05]. The tactile SES produced similar results, with significant main effects of timepoint [*F*(1,253) = 30.08, p < .001] and Gap versus No-Gap trial types [*F*(1,253) = 14.86, p < .001], but not frequency [*F*(3,251) = 2.31, p > .05], and no significant interaction [*F*(3,248) = 0.27, p > .05]. Critically, gap trials produced lower startle amplitudes relative to NG trials in both pre- and post-SS conditions for both SES types, indicating preserved gap detection despite overall amplitude increases.

## 4. Discussion

The GPIAS paradigm, first described by Turner et al. (2006), has become a widely adopted behavioral method for screening tinnitus in rodents. In this approach, a brief silent gap embedded within a continuous narrowband noise serves as the prepulse, and the carrier frequency is typically chosen to approximate the animal’s putative tinnitus pitch. Under normal hearing conditions, the presence of a gap suppresses the startle response, producing a lower ratio. In contrast, animals experiencing tinnitus exhibit reduced gap-induced inhibition - presumably because the tinnitus percept “fills in” the gap - resulting in larger startle amplitudes and an increased G:NG. Compared to operant conditioning paradigms such as active avoidance, go/no-go discrimination (Zuo et al., 2017), and two-alternative forced choice tasks (Hayes et al., 2023), GPIAS offers several practical advantages. Because it is based on an unconditioned reflex, it requires no pre-training, which not only reduces the time and resources needed to prepare animals for testing but also avoids the induction of conditioning-related neuroplastic changes in auditory processing that could confound the interpretation of tinnitus-related findings (Galazyuk & Hébert, 2015). Operant paradigms additionally carry the risk that animals may forget conditioned responses during extended tinnitus induction periods, which may reduce the reliability of longitudinal comparisons (Fabrizio-Stover et al., 2022). The reflexive nature of GPIAS also lends itself to high-throughput screening, allowing a large number of animals to be tested before and after tinnitus induction to separate tinnitus-positive from tinnitus-negative subjects (Galazyuk & Hébert, 2015).

Despite its widespread adoption, the validity of GPIAS as a tinnitus-specific assay remains a subject of ongoing debate. A central challenge is that the foundational “filling-in” hypothesis has not been consistently supported across studies. Human studies have reported GPIAS deficits that do not correspond to the subject’s matched tinnitus frequency, raising questions about whether reduced gap detection reflects tinnitus per se or broader changes in auditory temporal processing (Fournier & Hébert, 2013; Galazyuk & Hébert, 2015).

Compounding this, the lack of standardized criteria for defining a tinnitus-positive outcome across laboratories makes cross-study comparisons difficult, as current approaches often rely on simple population-level averaging of PPI values without consistent thresholds for classification (Schilling et al., 2017). A further source of variability is the floor effect identified by Lobarinas et al. (2013), who demonstrated that transient unilateral CHL induced by earplugging in rats significantly increased G:NG values - not due to tinnitus, but due to a disproportionate reduction in the NG startle amplitude that produced the appearance of reduced gap inhibition. Together, these issues highlight the need for methodological refinements that improve the specificity and reliability of GPIAS as a tinnitus screening tool. The present study sought to address several of these concerns directly: by employing a machine learning classifier to improve the accuracy of startle classification, comparing acoustic and tactile startle elicitors to disentangle hearing loss-related artifacts from true tinnitus-related gap detection deficits, and validating the paradigm across two distinct hearing impairment methods in CBA mice.

In experiment 1, startle amplitude input/output functions revealed a reduction in startle amplitude for both acoustic and tactile SES types following earplug insertion (Figure 3). The reduction observed during tactile SES testing likely reflects attenuation of the airpuff’s acoustic component by the earplug (Taylor, 1991). Contrary to expectations based on the rat study, we observed significant G:NG reductions across all carrier frequencies for the acoustic SES and at 16 and 20 kHz for the tactile SES. Examination of the startle amplitudes for gap and NG trials showed decreased startle responses during gap trials across all frequencies and both SES types, under earplug conditions. Together, these results suggest that unilateral CHL in CBA mice produces a generalized suppression of startle reactivity rather than a tinnitus-like deficit in gap detection, a pattern that is qualitatively distinct from the false-positive outcomes reported in rats.

In the second experiment, we used peripheral injections of SS to establish a positive control for tinnitus detection in the mouse model. After SS treatment, acoustic SES produced a significantly elevated startle input/output function, across all SES levels (Figure 5A), consistent with hyperacusis, a commonly reported effect of SS (Chen et al., 2014; Chen et al., 2024; Sun et al., 2009). A similar hyperresponsiveness was observed with tactile SES, again, attributable to SS effects on the acoustic component of the airpuff stimulus. In the MFG tests, both acoustic and tactile SES produced increased startle amplitudes in both gap and NG trials, resulting in inconsistent G:NG ratios that did not differ reliably across timepoints or frequencies; however, a significant interaction between frequency and timepoint was observed in the acoustic trials. Overall, despite the general SS-induced startle amplitude increases, amplitudes in gap trials were consistently smaller than NG trials for both SES types, indicating preserved gap inhibition. Therefore, the startle amplitude elevations appear to reflect a global hyperacusic response rather than a specific effect on gap processing. The multi-frequency assessment supports the value of using multiple carrier frequencies rather than relying on a single frequency, as frequency-specific effects on G:NG - such as the interaction observed for the acoustic SES - may be missed.

A key methodological distinction between the present study and Lobarinas et al. (2013) concerns how startle responses were defined and quantified. In that study, startle amplitude was measured using the RMS value from pre- and post-startle windows, an approach that does not distinguish true startle responses from low-amplitude noise. In contrast, our study employed a machine learning algorithm to identify and exclude non-startle responses, thereby minimizing contamination of the dataset. This distinction is likely consequential: if low-amplitude non-startle events were averaged together with inhibited startles from gap trials, baseline gap amplitudes may have been artificially reduced, lowering baseline G:NG values and exaggerating post-earplug increases to produce the appearance of a tinnitus-like effect. In our data, startle amplitudes decreased systematically across both gap and NG trials following CHL induction, consistent with reduced auditory responsivity. Because gap trials were already inhibited relative to NG trials at baseline, the proportional decrease in amplitude was greater for Gap trials, resulting in decreased rather than increased G:NG values after earplug placement. These findings suggest that differences in analytical methodology - specifically, how startle responses are defined and averaged - may account for the contrasting results between studies, and underscore the value of validated, automated classification methods for improving the reliability of GPIAS-based tinnitus screening.

Taken together, the frequency-dependent G:NG changes observed across carrier frequencies in experiment 2 support the utility of a multi-frequency screening approach, as these patterns are consistent with a frequency-specific ’filling-in’ effect tied to the animal’s putative tinnitus percept. This has important implications for future preclinical research, offering a more nuanced method for detecting tinnitus beyond traditional behavioral paradigms such as active avoidance, go/no-go discrimination, and two-alternative forced choice tasks, which rely on conditioned responses and take considerably longer to complete. However, the interpretation of these results is limited by several factors. The acoustic component of the airpuff introduces an auditory confound that cannot be fully controlled in the present design; a future experiment utilizing a silenced airpuff could prove useful in disentangling the role of sensorimotor gating from auditory sensitivity, particularly in hearing-impaired mice. Beyond this, a further limitation is the inherent difficulty in making direct comparisons between mouse and rat data. Mice and rats differ in baseline startle reactivity, audiometric range, and susceptibility to pharmacological and mechanical hearing manipulations, any of which could contribute to the divergent G:NG patterns observed across species. In particular, themouse’s extended high-frequency hearing range may alter how conductive hearing loss and salicylate-induced tinnitus are expressed behaviorally relative to the rat. Because analytical methodology and species were not independently varied across studies, it remains difficult to fully dissociate the contributions of biological and methodological factors to the observed differences in outcomes. Individual variability in salicylate response and earplug fit may have also contributed to within-group variance in G:NG values. Future studies should aim to replicate these findings across a broader range of frequencies, utilize a more precisely controlled tactile stimulus, and systematically vary analytical methodology within a single species to more directly test the contribution of classification approach to GPIAS outcomes, ultimately advancing the clinical utility of GPIAS as a preclinical tinnitus screening tool.

## CRediT authorship contribution statement

**Alyssa B. Price:** Data Curation, Formal Analysis, Investigation, Software, Visualization, Writing – original draft, Writing – review and editing. **Dimitri L. Brunelle:** Data Curation, Formal Analysis, Investigation, Software, Validation, Visualization, Writing – original draft, Writing – review and editing. **Collin R. Park:** Conceptualization, Formal Analysis, Methodology, Project Administration, Resources, Software, Supervision, Validation, Visualization, Writing – original draft, Writing – review and editing. **Andrea S. Lowe:** Conceptualization, Data Curation, Investigation, Methodology. **Edward Lobarinas:** Conceptualization, Writing – review and editing. **Joseph P. Walton:** Conceptualization, Funding Acquisition, Methodology, Project Administration, Resources, Supervision, Validation, Writing – review and editing.

## Acknowledgements

We would like to thank Dr. Elliott Brecht for his contributions to data collection. We would also like to thank our talented Undergraduate Research Assistants within the Global Center for Hearing and Speech Research for their contributions to data collection.

## Data Availability

Data will be made available on request.

## Glossary

GPIAS: Gap Prepulse Inhibition of the Acoustic Startle Reflex
ASR: Acoustic Startle Reflex
SES: Startle Elicitor Stimulus
PPI: Prepulse Inhibition
G:NG: Gap:No-Gap Ratio
SS: Sodium Salicylate
CHL: Conductive Hearing Loss
ABR: Auditory Brainstem Response
SIO: Startle Input/Output
ANOVA: Analysis of Variance
MFG: Multi-Frequency Gap
RMS: Root Mean Squared

---

**Figure S1.**
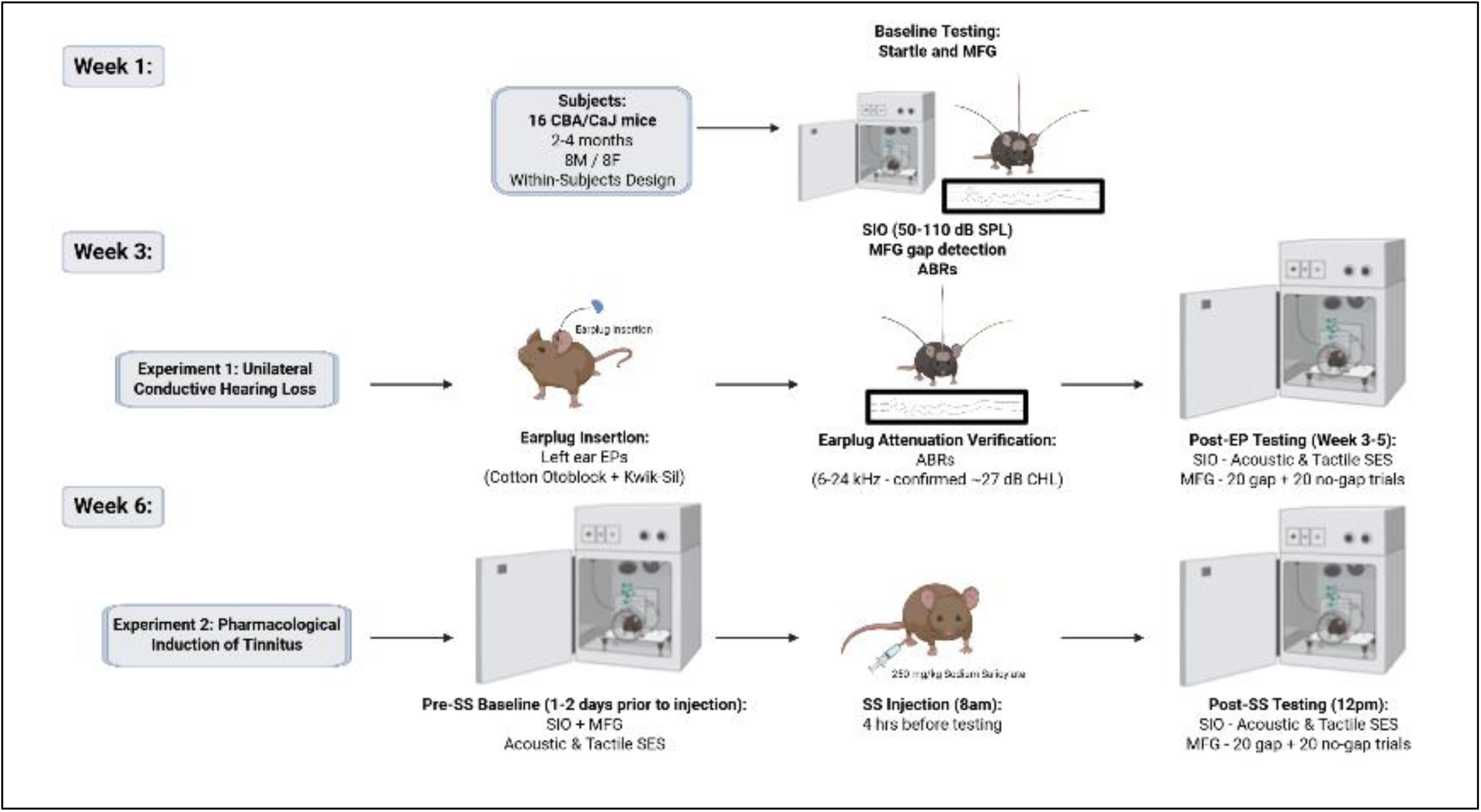
Methods Schematic for all Testing Procedures and Timelines.

**Figure S2.**
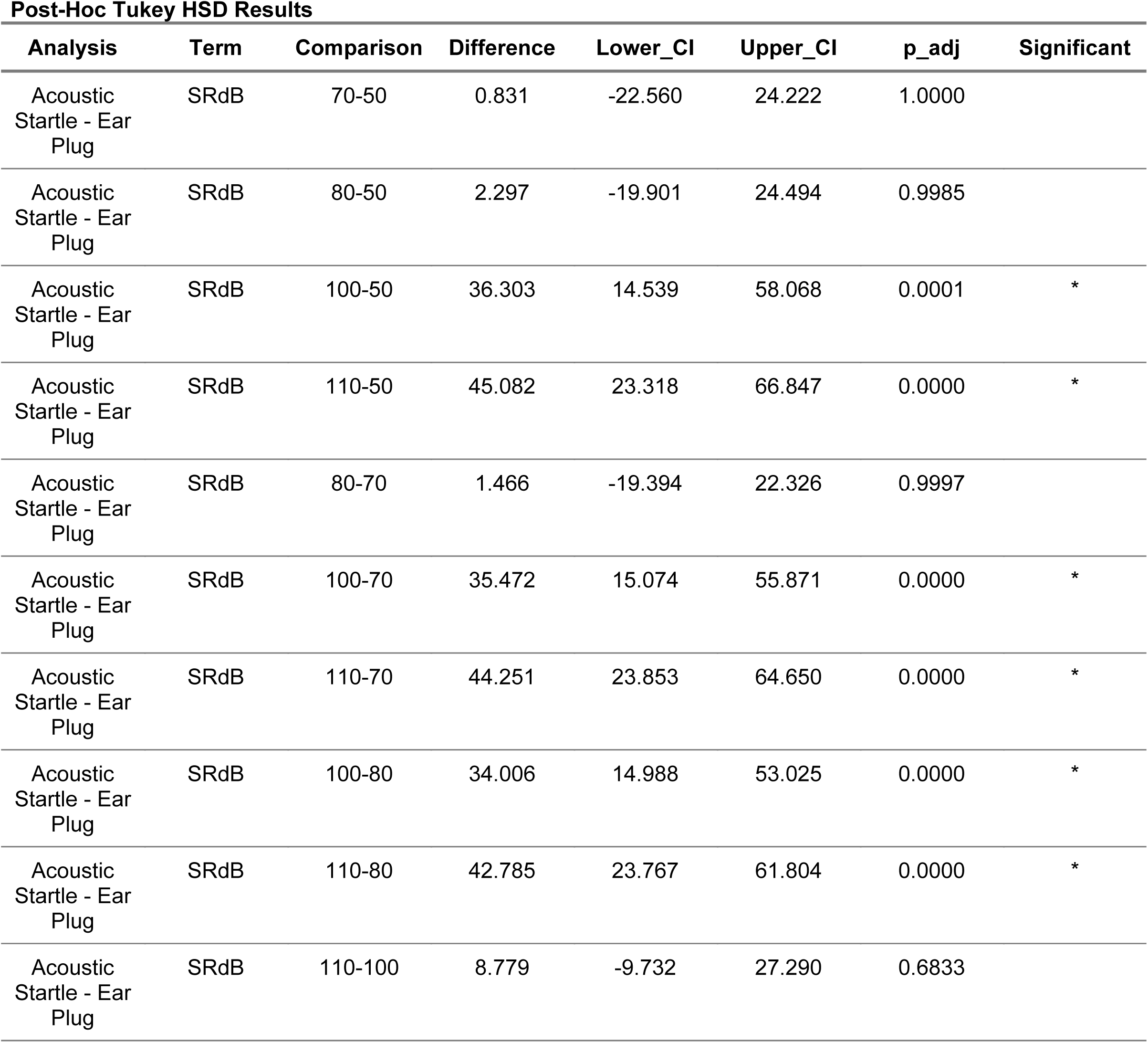
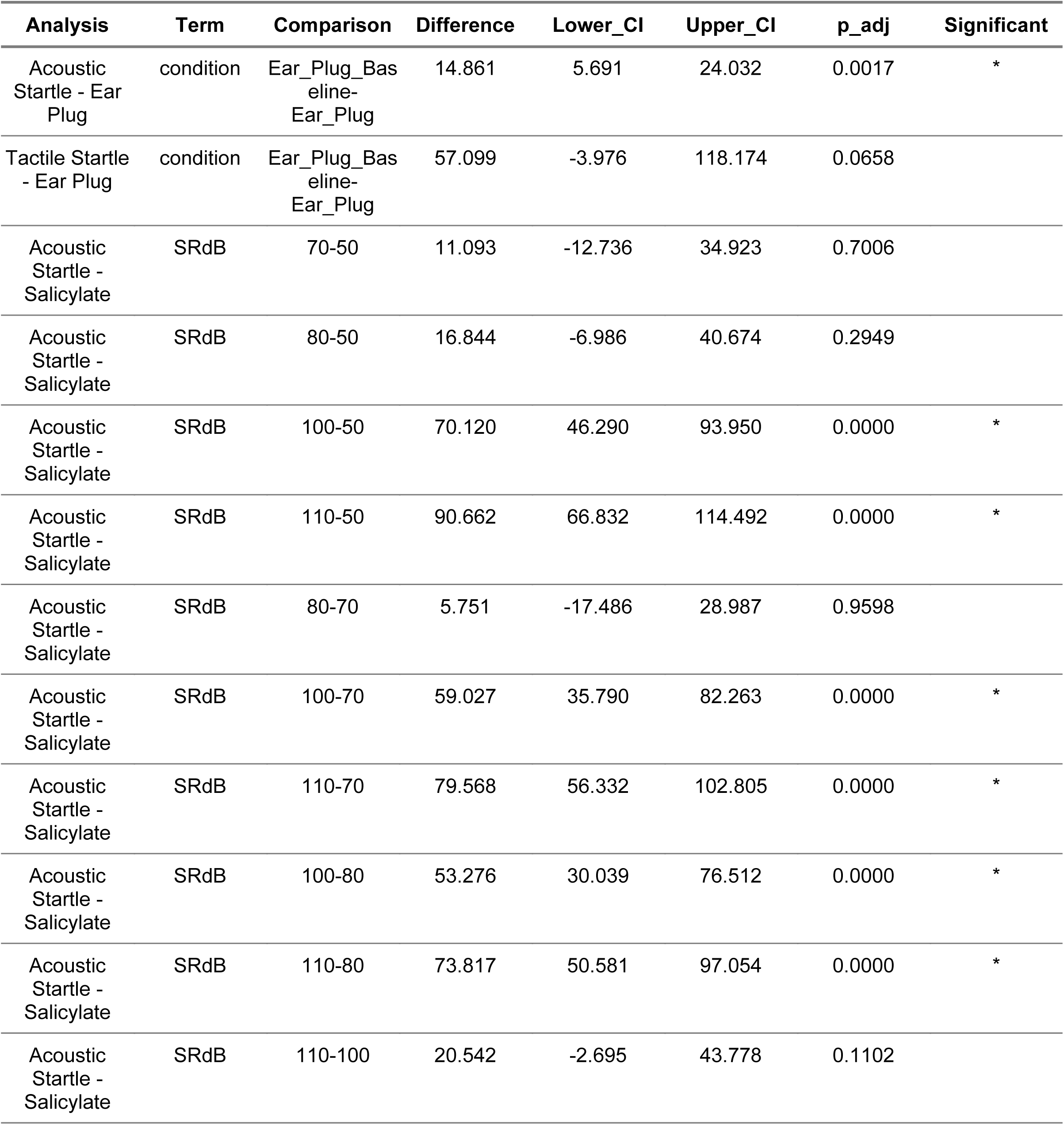
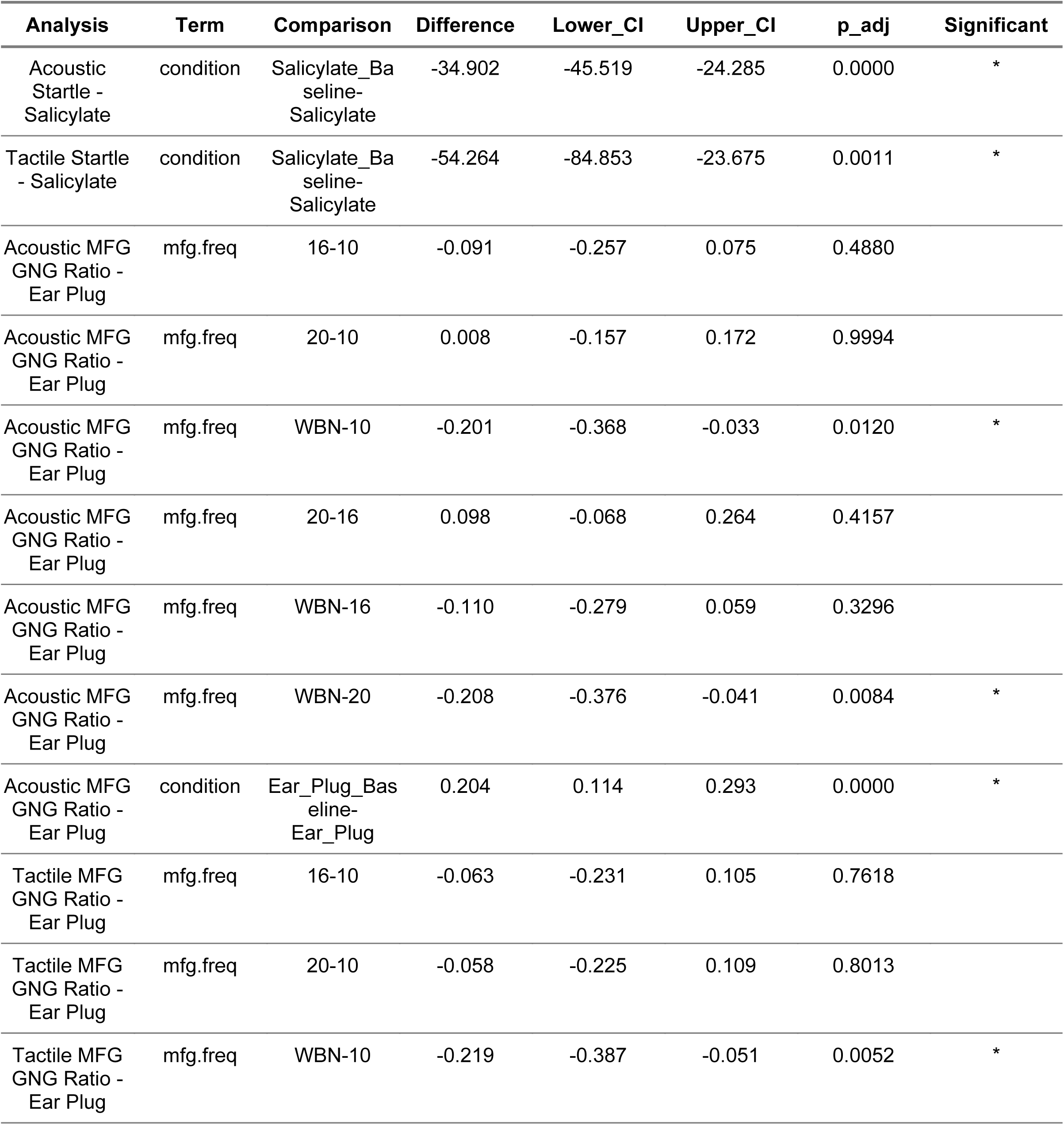
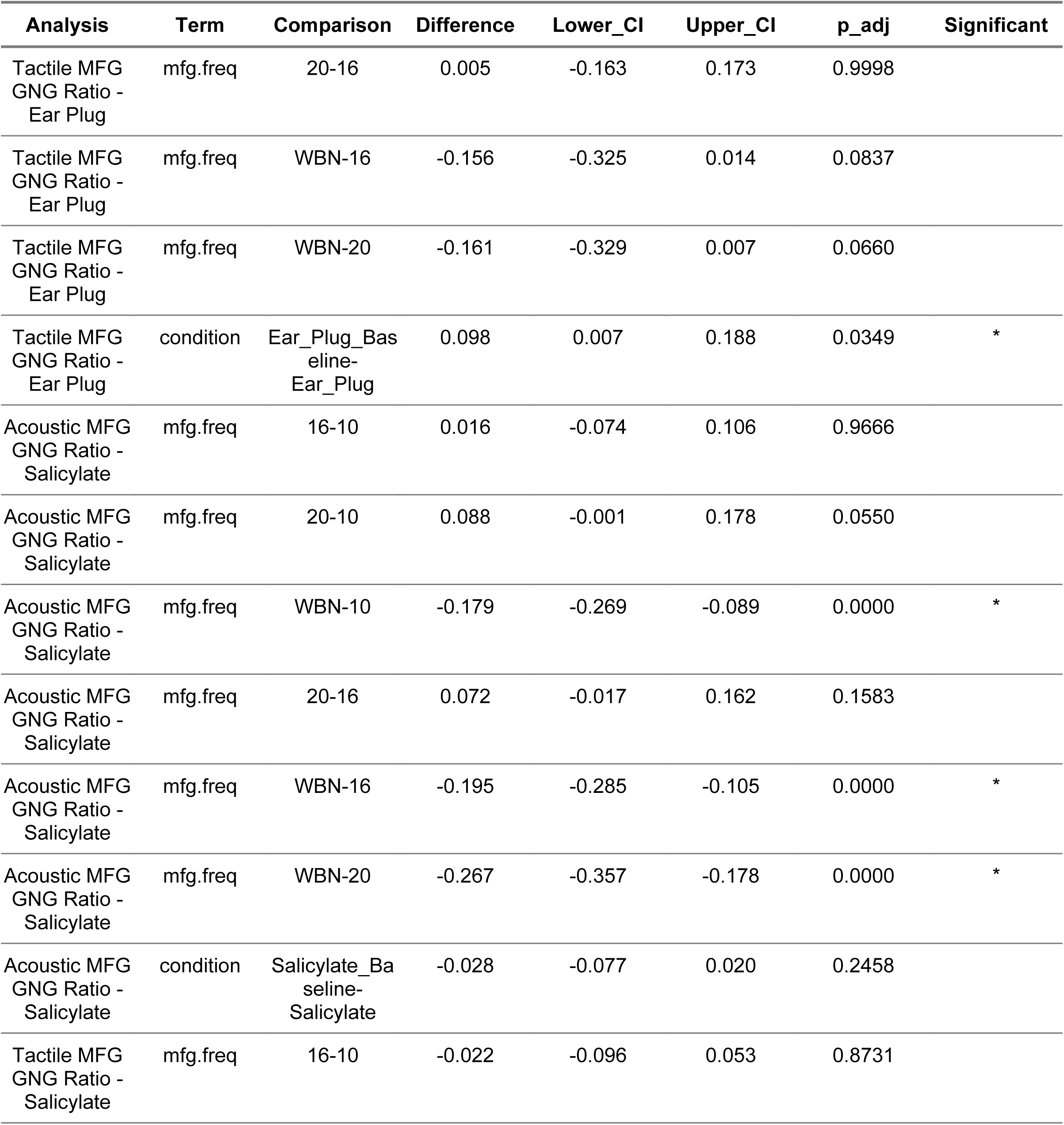
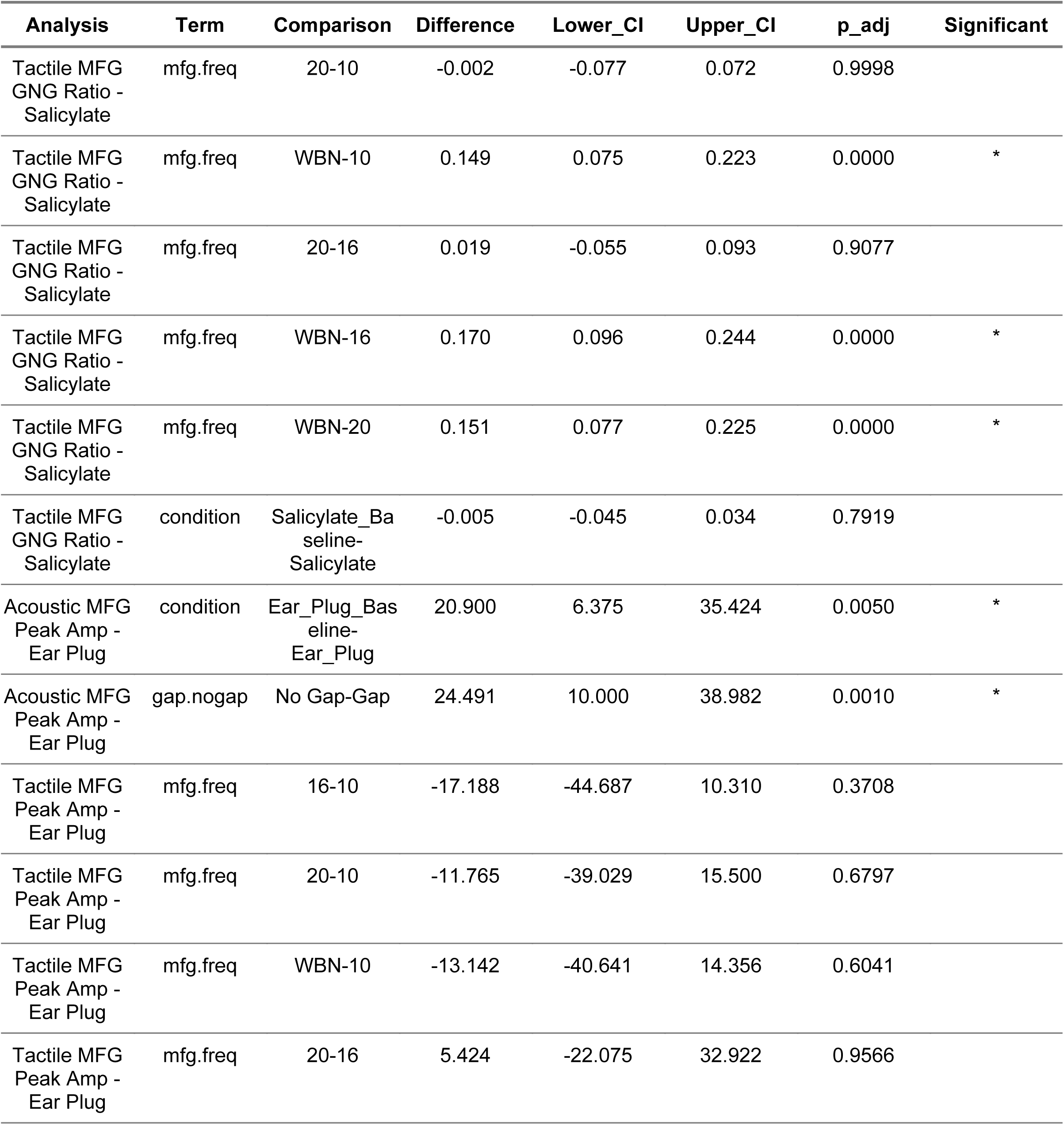
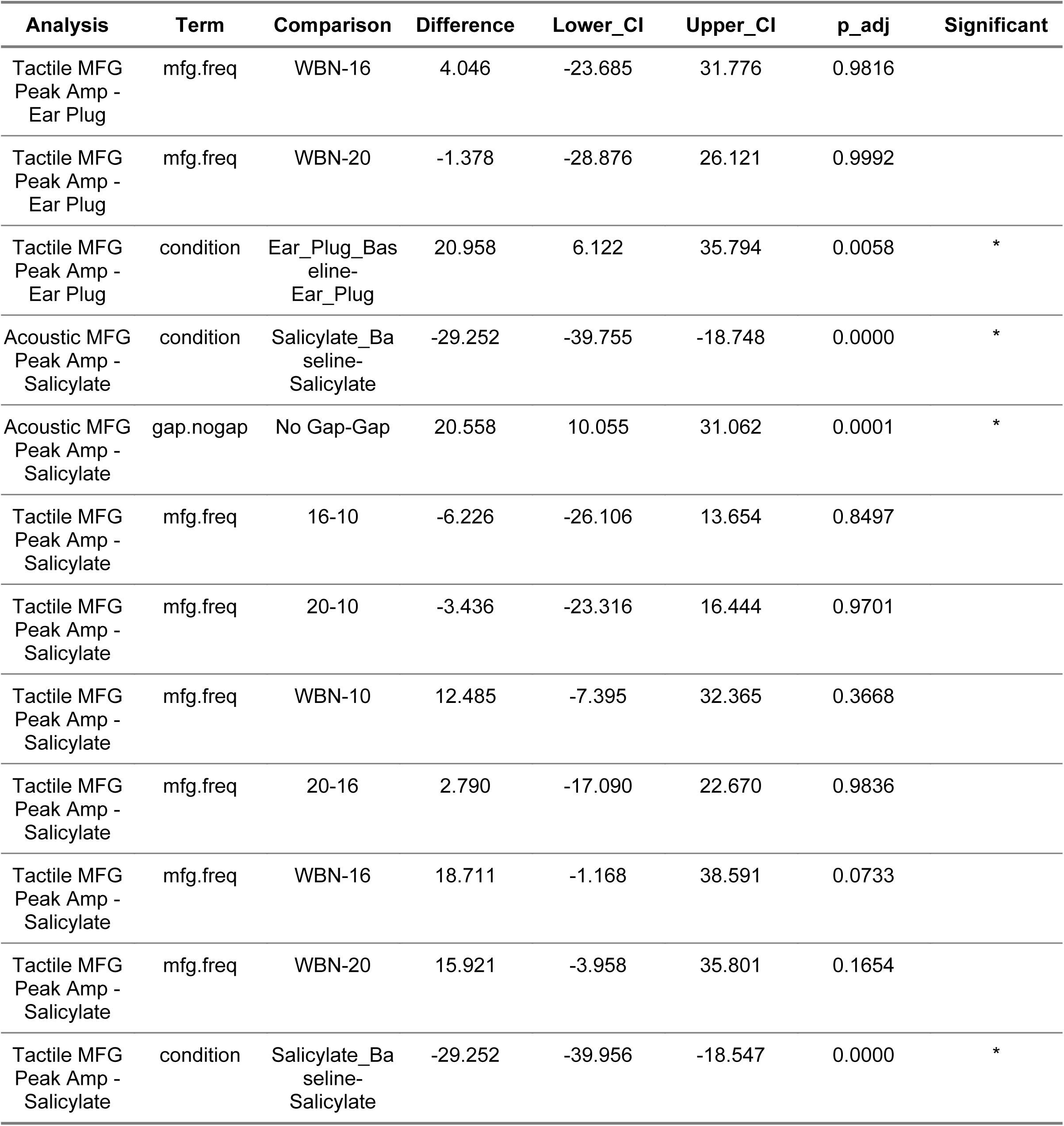
Post-Hoc Results Table.

## References

1. Barsz, K., Ison, J. R., Snell, K. B., & Walton, J. P. (2002). Behavioral and neural measures of auditory temporal acuity in aging humans and mice. Neurobiology of Aging, 23(4), 565–578. doi:10.1016/S0197-4580(02)00008-8

2. Bauer, C. A., & Brozoski, T. J. (2001). Assessing tinnitus and prospective tinnitus therapeutics using a psychophysical animal model. Journal of the Association for Research in Otolaryngology, 2(1), 54–64. doi:10.1007/s101620010030

3. Brecht, E.J., Scott, L.L., Ding, B., Zhu, X., Walton, J.P., 2022. A BK channel-targeted peptide induces age-dependent improvement in behavioral and neural sound representation. Neurobiol. Aging 110, 61–72. 10.1016/j.neurobiolaging.2021.10.014.

4. Brunelle, D.L., Park, C.R., Fawcett, T.J., Walton, J.P., 2025. Signal-in-noise detection across the lifespan in a mouse model of presbycusis. Hear. Res. 455, 109153. 10.1016/j.heares.2024.109153.

5. Chen, G., Radziwon, K. E., Kashanian, N., Manohar, S., & Salvi, R. (2014). Salicylate-induced auditory perceptual disorders and plastic changes in nonclassical auditory centers in rats. Neural Plasticity, 2014, 658741. doi:10.1155/2014/658741

6. Chen, J., Wang, X., Li, Z., Yuan, H., Wang, X., Yun, Y., et al. (2024). Thalamo-cortical neural mechanism of sodium salicylate-induced hyperacusis and anxiety-like behaviors. Communications Biology, 7(1), 1–17. doi:10.1038/s42003-024-07040-5

7. Davis, M. (1984). The mammalian startle response. In R. C. Eaton (Ed.), Neural mechanisms of startle behavior (pp. 287–351). Boston, MA: Springer US. Retrieved from 10.1007/978-1-4899-2286-1_10

8. Dawson, M. E., Schell, A. M., & Böhmelt, A. H. (Eds.). (1999). Startle modification: Implications for neuroscience, cognitive science, and clinical science. New York, NY: Cambridge University Press.

9. De Ridder, D., Schlee, W., Vanneste, S., Londero, A., Weisz, N., Kleinjung, T., et al. (2021). Tinnitus and tinnitus disorder: Theoretical and operational definitions (an international multidisciplinary proposal). Progress in Brain Research, 260, 1–25. doi:10.1016/bs.pbr.2020.12.002

10. Eggermont, J. J., & Roberts, L. E. (2015). Tinnitus: Animal models and findings in humans. Cell and Tissue Research, 361(1), 311–336. doi:10.1007/s00441-014-1992-8

11. Fabrizio-Stover, E. M., Nichols, G., Corcoran, J., Jain, A., Burghard, A. L., Lee, C. M., & Oliver, D. L. (2022). Comparison of two behavioral tests for tinnitus assessment in mice. Frontiers in Behavioral Neuroscience, 16, 995422. doi:10.3389/fnbeh.2022.995422

12. Fawcett, T. J., Cooper, C. S., Longenecker, R. J., & Walton, J. P. (2020). Automated classification of acoustic startle reflex waveforms in young CBA/CaJ mice using machine learning. Journal of Neuroscience Methods, 344, 108853. doi:10.1016/j.jneumeth.2020.108853

13. Fawcett, T. J., Cooper, C. S., Longenecker, R. J., & Walton, J. P. (2021). Machine learning, waveform preprocessing and feature extraction methods for classification of acoustic startle waveforms. MethodsX, 8, 101166. doi:10.1016/j.mex.2020.101166

14. Fawcett, T. J., Longenecker, R. J., Brunelle, D. L., Berger, J. I., Wallace, M. N., Galazyuk, A. V., et al. (2023). Universal automated classification of the acoustic startle reflex using machine learning. Hearing Research, 428, 108667. doi:10.1016/j.heares.2022.108667

15. Fournier, P., & Hébert, S. (2013). Gap detection deficits in humans with tinnitus as assessed with the acoustic startle paradigm: Does tinnitus fill in the gap? Hearing Research, 295, 16–23. doi:10.1016/j.heares.2012.05.011

16. Galazyuk, A., & Hébert, S. (2015). Gap-prepulse inhibition of the acoustic startle reflex (GPIAS) for tinnitus assessment: Current status and future directions. Frontiers in Neurology, 6, 88. doi:10.3389/fneur.2015.00088

17. Gao, J., Wu, X., & Zuo, J. (2004). Targeting hearing genes in mice. Brain Research. Molecular Brain Research, 132(2), 192–207. doi:10.1016/j.molbrainres.2004.06.035

18. García-Hernández, S., & Rubio, M. E. (2022). Role of GluA4 in the acoustic and tactile startle responses. Hearing Research, 414, 108410. doi:10.1016/j.heares.2021.108410

19. Grimsley, C. A., Longenecker, R. J., Rosen, M. J., Young, J. W., Grimsley, J. M., & Galazyuk, A. V. (2015). An improved approach to separating startle data from noise. Journal of Neuroscience Methods, 253, 206–217. doi:10.1016/j.jneumeth.2015.07.001

20. Halonen, J., Hinton, A.S., Frisina, R.D., Ding, B., Zhu, X., Walton, J.P., 2016. Long-term treatment with aldosterone slows the progression of age-related hearing loss. Hear. Res. 336, 63–71. 10.1016/j.heares.2016.05.001.

21. Hayes, S.H., Beh, K., Typlt, M., Schormans, A.L., Stolzberg, D., & Allman, B.L. (2023). Using an appetitive operant conditioning paradigm to screen rats for tinnitus induced by intense sound exposure: Experimental considerations and interpretation. Frontiers in Neuroscience, 17.

22. Henry, J. A., & Meikle, M. B. (2000). Psychoacoustic measures of tinnitus. Journal of the American Academy of Audiology, 11(3), 138–155. Retrieved from https://pubmed.ncbi.nlm.nih.gov/10755810/

23. Ison, J. R., McAdam, D. W., & Hammond, G. R. (1973). Latency and amplitude changes in the acoustic startle reflex of the rat produced by variation in auditory prestimulation. Physiology & Behavior, 10(6), 1035–1039. doi:10.1016/0031-9384(73)90185-6

24. Ison, J. R., O’Connor, K., Bowen, G. P., & Bocirnea, A. (1991). Temporal resolution of gaps in noise by the rat is lost with functional decortication. Behavioral Neuroscience, 105(1), 33–40. doi:10.1037/0735-7044.105.1.33

25. Jarach, C. M., Lugo, A., Scala, M., van den Brandt, P. A., Cederroth, C. R., Odone, A., et al. (2022). Global prevalence and incidence of tinnitus: A systematic review and meta-analysis. JAMA Neurology, 79(9), 888– 900. doi:10.1001/jamaneurol.2022.2189

26. Landis, C., & Hunt, W. (1939). The startle pattern. Oxford, England: Farrar & Rinehart. Retrieved from https://psycnet.apa.org/record/1939-03049-000

27. Lobarinas, E., Hayes, S. H., & Allman, B. L. (2013). The gap-startle paradigm for tinnitus screening in animal models: Limitations and optimization. Hearing Research, 295, 150–160. doi:10.1016/j.heares.2012.06.001

28. Longenecker, R. J., & Galazyuk, A. V. (2012). Methodological optimization of tinnitus assessment using prepulse inhibition of the acoustic startle reflex. Brain Research, 1485, 54–62. doi:10.1016/j.brainres.2012.02.067

29. Longenecker, R. J., & Galazyuk, A. V. (2011). Development of tinnitus in CBA/CaJ mice following sound exposure. Journal of the Association for Research in Otolaryngology: JARO, 12(5), 647–658. doi:10.1007/s10162-011-0276-1

30. Lowe, A. S., & Walton, J. P. (2015). Alterations in peripheral and central components of the auditory brainstem response: A neural assay of tinnitus. Plos One, 10(2), e0117228. doi:10.1371/journal.pone.0117228

31. Plappert, C., Pilz, P., & Schnitzler, H. (2004). Factors governing prepulse inhibition and prepulse facilitation of the acoustic startle response in mice. Behavioural Brain Research, 152, 403–12. doi:10.1016/j.bbr.2003.10.025

32. Radziwon, K., Holfoth, D., Lindner, J., Kaier-Green, Z., Bowler, R., Urban, M., et al. (2017). Salicylate-induced hyperacusis in rats: Dose-and frequency-dependent effects. Hearing Research, 350, 133–138. doi:10.1016/j.heares.2017.04.004

33. Refat, F., Wertz, J., Hinrichs, P., Klose, U., Samy, H., Abdelkader, R. M., Saemisch, J., Hofmeier, B., Singer, W., Rüttiger, L., Knipper, M., & Wolpert, S. (2021). Co-occurrence of hyperacusis accelerates with tinnitus burden over time and requires medical care. Frontiers in Neurology, 12, 627522. doi:10.3389/fneur.2021.627522

34. Schecklmann, M., Landgrebe, M., & Langguth, B. (2014). Phenotypic characteristics of hyperacusis in tinnitus. PLOS ONE, 9(1), e86944. doi:10.1371/journal.pone.0086944

35. Schilling, A., Krauss, P., Gerum, R., Metzner, C., Tziridis, K., & Schulze, H. (2017). A new statistical approach for the evaluation of gap-prepulse inhibition of the acoustic startle reflex (GPIAS) for tinnitus assessment. Frontiers in Behavioral Neuroscience, 11, 198. doi:10.3389/fnbeh.2017.00198

36. Sun, W., Lu, J., Stolzberg, D., Gray, L., Deng, A., Lobarinas, E., et al. (2009). Salicylate increases the gain of the central auditory system. Neuroscience, 159(1), 325–334. doi:10.1016/j.neuroscience.2008.12.024

37. Taylor, B. K., Casto, R., & Printz, M. P. (1991). Dissociation of tactile and acoustic components in air puff startle. Physiology & Behavior, 49(3), 527–532. doi:10.1016/0031-9384(91)90275-S

38. Turner, J. G., Brozoski, T. J., Bauer, C. A., Parrish, J. L., Myers, K., Hughes, L. F., et al. (2006). Gap detection deficits in rats with tinnitus: A potential novel screening tool. Behavioral Neuroscience, 120(1), 188–195. doi:10.1037/0735-7044.120.1.188

39. Watts, E. J., Fackrell, K., Smith, S., Sheldrake, J., Haider, H., & Hoare, D. J. (2018). Why is tinnitus a problem? A qualitative analysis of problems reported by tinnitus patients. Trends in Hearing, 22, 2331216518812250. doi:10.1177/2331216518812250

40. Yang, G., Lobarinas, E., Zhang, L., Turner, J., Stolzberg, D., Salvi, R., et al. (2007). Salicylate induced tinnitus: Behavioral measures and neural activity in auditory cortex of awake rats. Hearing Research, 226(1-2), 244–253. doi:10.1016/j.heares.2006.06.013

