## Supplementary material for "Validating the Gap-Startle Paradigm for Tinnitus Detection: A Machine Learning Approach in CBA/CaJ Mice": S1

### Post-Hoc Tukey HSD Results

| **Analysis** | **Term** | **Comparison** | **Difference** | **Lower_CI** | **Upper_CI** | **p_adj** | **Significant** |
| --- | --- | --- | --- | --- | --- | --- | --- |
| Acoustic Startle - Ear Plug | SRdB | 70-50 | 0.831 | -22.560 | 24.222 | 1.0000 |  |
| Acoustic Startle - Ear Plug | SRdB | 80-50 | 2.297 | -19.901 | 24.494 | 0.9985 |  |
| Acoustic Startle - Ear Plug | SRdB | 100-50 | 36.303 | 14.539 | 58.068 | 0.0001 | * |
| Acoustic Startle - Ear Plug | SRdB | 110-50 | 45.082 | 23.318 | 66.847 | 0.0000 | * |
| Acoustic Startle - Ear Plug | SRdB | 80-70 | 1.466 | -19.394 | 22.326 | 0.9997 |  |
| Acoustic Startle - Ear Plug | SRdB | 100-70 | 35.472 | 15.074 | 55.871 | 0.0000 | * |
| Acoustic Startle - Ear Plug | SRdB | 110-70 | 44.251 | 23.853 | 64.650 | 0.0000 | * |
| Acoustic Startle - Ear Plug | SRdB | 100-80 | 34.006 | 14.988 | 53.025 | 0.0000 | * |
| Acoustic Startle - Ear Plug | SRdB | 110-80 | 42.785 | 23.767 | 61.804 | 0.0000 | * |
| Acoustic Startle - Ear Plug | SRdB | 110-100 | 8.779 | -9.732 | 27.290 | 0.6833 |  |
| Acoustic Startle - Ear Plug | condition | Ear_Plug_Baseline-Ear_Plug | 14.861 | 5.691 | 24.032 | 0.0017 | * |
| Tactile Startle - Ear Plug | condition | Ear_Plug_Baseline-Ear_Plug | 57.099 | -3.976 | 118.174 | 0.0658 |  |
| Acoustic Startle - Salicylate | SRdB | 70-50 | 11.093 | -12.736 | 34.923 | 0.7006 |  |
| Acoustic Startle - Salicylate | SRdB | 80-50 | 16.844 | -6.986 | 40.674 | 0.2949 |  |
| Acoustic Startle - Salicylate | SRdB | 100-50 | 70.120 | 46.290 | 93.950 | 0.0000 | * |
| Acoustic Startle - Salicylate | SRdB | 110-50 | 90.662 | 66.832 | 114.492 | 0.0000 | * |
| Acoustic Startle - Salicylate | SRdB | 80-70 | 5.751 | -17.486 | 28.987 | 0.9598 |  |
| Acoustic Startle - Salicylate | SRdB | 100-70 | 59.027 | 35.790 | 82.263 | 0.0000 | * |
| Acoustic Startle - Salicylate | SRdB | 110-70 | 79.568 | 56.332 | 102.805 | 0.0000 | * |
| Acoustic Startle - Salicylate | SRdB | 100-80 | 53.276 | 30.039 | 76.512 | 0.0000 | * |
| Acoustic Startle - Salicylate | SRdB | 110-80 | 73.817 | 50.581 | 97.054 | 0.0000 | * |
| Acoustic Startle - Salicylate | SRdB | 110-100 | 20.542 | -2.695 | 43.778 | 0.1102 |  |
| Acoustic Startle - Salicylate | condition | Salicylate_Baseline-Salicylate | -34.902 | -45.519 | -24.285 | 0.0000 | * |
| Tactile Startle - Salicylate | condition | Salicylate_Baseline-Salicylate | -54.264 | -84.853 | -23.675 | 0.0011 | * |
| Acoustic MFG GNG Ratio - Ear Plug | mfg.freq | 16-10 | -0.091 | -0.257 | 0.075 | 0.4880 |  |
| Acoustic MFG GNG Ratio - Ear Plug | mfg.freq | 20-10 | 0.008 | -0.157 | 0.172 | 0.9994 |  |
| Acoustic MFG GNG Ratio - Ear Plug | mfg.freq | WBN-10 | -0.201 | -0.368 | -0.033 | 0.0120 | * |
| Acoustic MFG GNG Ratio - Ear Plug | mfg.freq | 20-16 | 0.098 | -0.068 | 0.264 | 0.4157 |  |
| Acoustic MFG GNG Ratio - Ear Plug | mfg.freq | WBN-16 | -0.110 | -0.279 | 0.059 | 0.3296 |  |
| Acoustic MFG GNG Ratio - Ear Plug | mfg.freq | WBN-20 | -0.208 | -0.376 | -0.041 | 0.0084 | * |
| Acoustic MFG GNG Ratio - Ear Plug | condition | Ear_Plug_Baseline-Ear_Plug | 0.204 | 0.114 | 0.293 | 0.0000 | * |
| Tactile MFG GNG Ratio - Ear Plug | mfg.freq | 16-10 | -0.063 | -0.231 | 0.105 | 0.7618 |  |
| Tactile MFG GNG Ratio - Ear Plug | mfg.freq | 20-10 | -0.058 | -0.225 | 0.109 | 0.8013 |  |
| Tactile MFG GNG Ratio - Ear Plug | mfg.freq | WBN-10 | -0.219 | -0.387 | -0.051 | 0.0052 | * |
| Tactile MFG GNG Ratio - Ear Plug | mfg.freq | 20-16 | 0.005 | -0.163 | 0.173 | 0.9998 |  |
| Tactile MFG GNG Ratio - Ear Plug | mfg.freq | WBN-16 | -0.156 | -0.325 | 0.014 | 0.0837 |  |
| Tactile MFG GNG Ratio - Ear Plug | mfg.freq | WBN-20 | -0.161 | -0.329 | 0.007 | 0.0660 |  |
| Tactile MFG GNG Ratio - Ear Plug | condition | Ear_Plug_Baseline-Ear_Plug | 0.098 | 0.007 | 0.188 | 0.0349 | * |
| Acoustic MFG GNG Ratio - Salicylate | mfg.freq | 16-10 | 0.016 | -0.074 | 0.106 | 0.9666 |  |
| Acoustic MFG GNG Ratio - Salicylate | mfg.freq | 20-10 | 0.088 | -0.001 | 0.178 | 0.0550 |  |
| Acoustic MFG GNG Ratio - Salicylate | mfg.freq | WBN-10 | -0.179 | -0.269 | -0.089 | 0.0000 | * |
| Acoustic MFG GNG Ratio - Salicylate | mfg.freq | 20-16 | 0.072 | -0.017 | 0.162 | 0.1583 |  |
| Acoustic MFG GNG Ratio - Salicylate | mfg.freq | WBN-16 | -0.195 | -0.285 | -0.105 | 0.0000 | * |
| Acoustic MFG GNG Ratio - Salicylate | mfg.freq | WBN-20 | -0.267 | -0.357 | -0.178 | 0.0000 | * |
| Acoustic MFG GNG Ratio - Salicylate | condition | Salicylate_Baseline-Salicylate | -0.028 | -0.077 | 0.020 | 0.2458 |  |
| Tactile MFG GNG Ratio - Salicylate | mfg.freq | 16-10 | -0.022 | -0.096 | 0.053 | 0.8731 |  |
| Tactile MFG GNG Ratio - Salicylate | mfg.freq | 20-10 | -0.002 | -0.077 | 0.072 | 0.9998 |  |
| Tactile MFG GNG Ratio - Salicylate | mfg.freq | WBN-10 | 0.149 | 0.075 | 0.223 | 0.0000 | * |
| Tactile MFG GNG Ratio - Salicylate | mfg.freq | 20-16 | 0.019 | -0.055 | 0.093 | 0.9077 |  |
| Tactile MFG GNG Ratio - Salicylate | mfg.freq | WBN-16 | 0.170 | 0.096 | 0.244 | 0.0000 | * |
| Tactile MFG GNG Ratio - Salicylate | mfg.freq | WBN-20 | 0.151 | 0.077 | 0.225 | 0.0000 | * |
| Tactile MFG GNG Ratio - Salicylate | condition | Salicylate_Baseline-Salicylate | -0.005 | -0.045 | 0.034 | 0.7919 |  |
| Acoustic MFG Peak Amp - Ear Plug | condition | Ear_Plug_Baseline-Ear_Plug | 20.900 | 6.375 | 35.424 | 0.0050 | * |
| Acoustic MFG Peak Amp - Ear Plug | gap.nogap | No Gap-Gap | 24.491 | 10.000 | 38.982 | 0.0010 | * |
| Tactile MFG Peak Amp - Ear Plug | mfg.freq | 16-10 | -17.188 | -44.687 | 10.310 | 0.3708 |  |
| Tactile MFG Peak Amp - Ear Plug | mfg.freq | 20-10 | -11.765 | -39.029 | 15.500 | 0.6797 |  |
| Tactile MFG Peak Amp - Ear Plug | mfg.freq | WBN-10 | -13.142 | -40.641 | 14.356 | 0.6041 |  |
| Tactile MFG Peak Amp - Ear Plug | mfg.freq | 20-16 | 5.424 | -22.075 | 32.922 | 0.9566 |  |
| Tactile MFG Peak Amp - Ear Plug | mfg.freq | WBN-16 | 4.046 | -23.685 | 31.776 | 0.9816 |  |
| Tactile MFG Peak Amp - Ear Plug | mfg.freq | WBN-20 | -1.378 | -28.876 | 26.121 | 0.9992 |  |
| Tactile MFG Peak Amp - Ear Plug | condition | Ear_Plug_Baseline-Ear_Plug | 20.958 | 6.122 | 35.794 | 0.0058 | * |
| Acoustic MFG Peak Amp - Salicylate | condition | Salicylate_Baseline-Salicylate | -29.252 | -39.755 | -18.748 | 0.0000 | * |
| Acoustic MFG Peak Amp - Salicylate | gap.nogap | No Gap-Gap | 20.558 | 10.055 | 31.062 | 0.0001 | * |
| Tactile MFG Peak Amp - Salicylate | mfg.freq | 16-10 | -6.226 | -26.106 | 13.654 | 0.8497 |  |
| Tactile MFG Peak Amp - Salicylate | mfg.freq | 20-10 | -3.436 | -23.316 | 16.444 | 0.9701 |  |
| Tactile MFG Peak Amp - Salicylate | mfg.freq | WBN-10 | 12.485 | -7.395 | 32.365 | 0.3668 |  |
| Tactile MFG Peak Amp - Salicylate | mfg.freq | 20-16 | 2.790 | -17.090 | 22.670 | 0.9836 |  |
| Tactile MFG Peak Amp - Salicylate | mfg.freq | WBN-16 | 18.711 | -1.168 | 38.591 | 0.0733 |  |
| Tactile MFG Peak Amp - Salicylate | mfg.freq | WBN-20 | 15.921 | -3.958 | 35.801 | 0.1654 |  |
| Tactile MFG Peak Amp - Salicylate | condition | Salicylate_Baseline-Salicylate | -29.252 | -39.956 | -18.547 | 0.0000 | * |
