## Supplementary figures and images for "Validating the Gap-Startle Paradigm for Tinnitus Detection: A Machine Learning Approach in CBA/CaJ Mice"

### S2

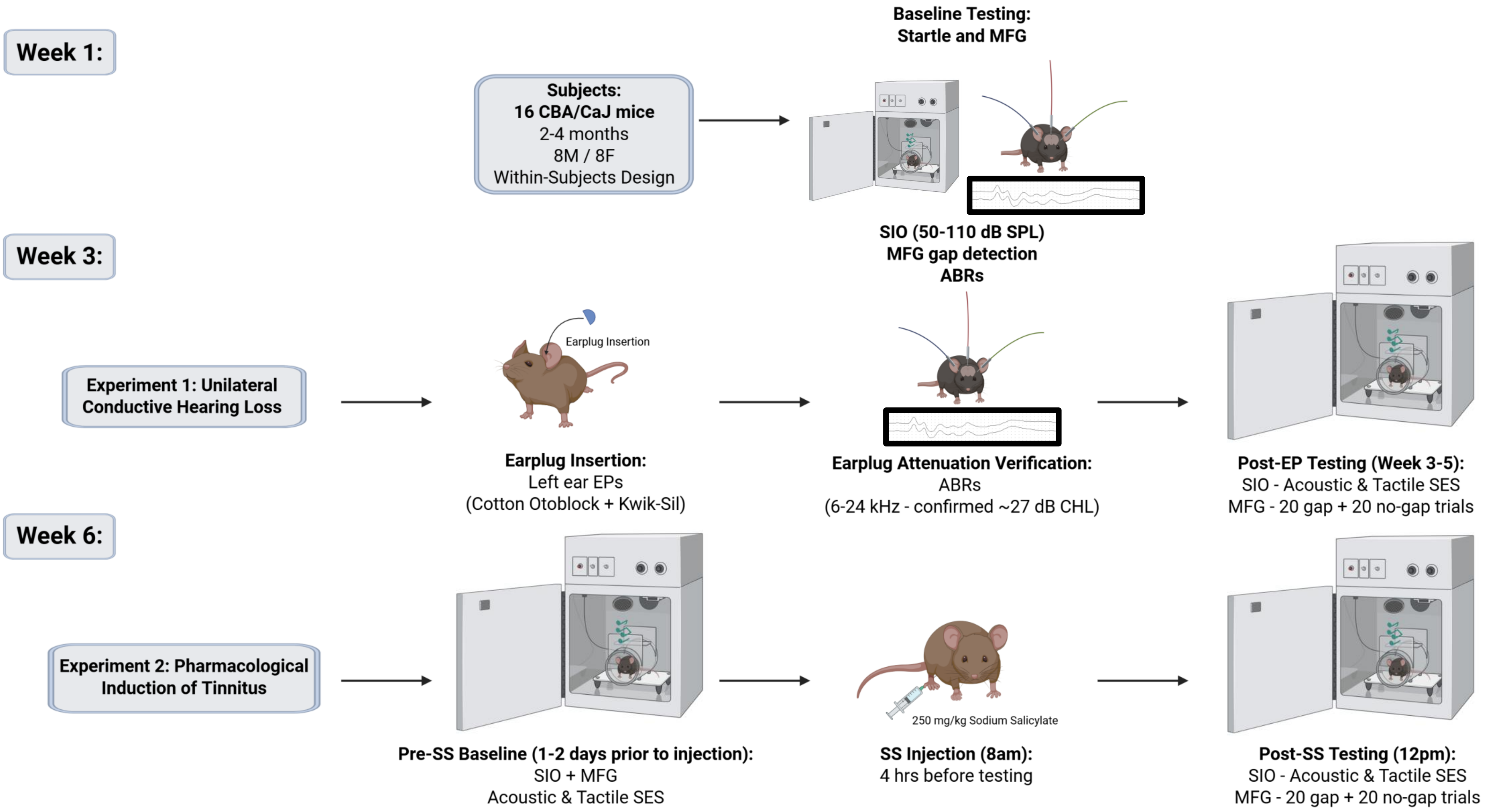
